# High-throughput neural stem cell-based drug screening identifies S6K1 inhibition as a selective vulnerability in SHH-medulloblastoma

**DOI:** 10.1101/2024.01.31.574335

**Authors:** Leilei Zhou, Niek van Bree, Lola Boutin, Simon Moussaud, Magdalena Otrocka, Anna Falk, Margareta Wilhelm

## Abstract

**Background:** Medulloblastoma (MB) is one of the most common malignant brain tumors in children. Current treatments have increased overall survival but can lead to devastating side effects and late complications in survivors, emphasizing the need for new, improved targeted therapies that specifically eliminate tumor cells while sparing the normally developing brain.

**Methods:** Here, we used a SHH-MB model based on a patient-derived neuroepithelial stem (NES) cell system for an unbiased high-throughput screen with a library of 172 compounds with known targets. Compounds were evaluated in both healthy neural stem cells and tumor cells derived from the same patient. Based on the difference of cell viability and drug sensitivity score between normal cells and tumor cells, hit compounds were selected and further validated *in vitro* and *in vivo*.

**Results:** We identified PF4708671 (S6K1 inhibitor) as a potential agent that selectively targets Sonic Hedgehog (SHH) driven MB tumor cells while sparing neural stem cells and differentiated neurons. Subsequent validation studies confirmed that PF4708671 inhibited the growth of SHH-MB tumor cells both *in vitro* and *in vivo*, and that knockdown of S6K1 resulted in reduced tumor formation.

**Conclusion:** Overall, our results suggest that inhibition of S6K1 specifically affects tumor growth, whereas it has less effect on non-tumor cells. Our data also show that the NES cell platform can be used to identify potentially effective new therapies and targets for SHH-MB.

**Key points:**

- High-throughput screening system using the NES model identifies efficient compounds and targets against SHH-MB.
- S6K1 inhibition shows selectivity toward tumor cells while having less effect on normal neural stem cells and neurons.

**Importance of the study:** Current treatment modalities for medulloblastoma have improved overall survival but also come with detrimental side effects for survivors. Therefore, novel treatment options need to be developed which will specifically target the tumor cells while sparing the healthy brain. In this study, we tested a library of compounds targeting commonly dysregulated oncogenic pathways on both normal neural stem cells and SHH-MB tumor cells derived from the same patients. Interestingly, we found that most compounds including commonly used targeted therapy such as PI3K or mTOR inhibition, albeit effective, affected tumor cells and normal cells similarly. However, inhibition of the downstream effector S6K1 preferentially targeted tumor cells both *in vitro* and *in vivo*. These results thus reveal potential targets for translational studies of novel therapies that specifically target medulloblastoma tumor cells.

## Introduction

Medulloblastoma (MB) is a malignant pediatric brain tumor which arises from a discrete population of neural stem or progenitor cells in the cerebellum or brainstem where the normal differentiation program has stalled [1, 2]. Multi-omics analyses divide MB into four consensus molecular subgroups, Wingless (WNT), Sonic hedgehog (SHH), Group 3 and Group 4 with different characteristics in terms of genetic and epigenetic changes, age of onset, and clinical outcome [1, 3]. SHH-MB is driven by pathogenic activation of the SHH pathway, which also drives normal proliferation of granule neural precursors (GNPs) during cerebellar development [4, 5]. The SHH subgroup accounts for 30% of MB cases and can occur in all age groups [6–9]. Gene expression and DNA methylation profiles further subdivide SHH-MB into four subtypes, SHH-1(−β), SHH-2(−γ), SHH-3(−α), and SHH-4(−δ) [10–12]. Inactivating mutations or loss of *PTCH1* are detected almost equally frequently in all SHH subtypes, whereas *MYCN* or *GLI2* amplifications and *TP53* mutations are more common in the SHH-3 subtype (children) [13]. *SMO* mutations are highly enriched in the SHH-4 subtype (adolescents and adults), while *SUFU* mutations are almost exclusively found in SHH-1 and SHH-2 (infants) [10, 11]. Among all SHH subtypes, SHH-1 has a higher incidence of metastasis, while SHH-2 and SHH-4 have a better survival rate [14].

Current standard treatment of MB includes surgery followed by intra-craniospinal irradiation and multidrug chemotherapy [1, 15]. Although the overall survival rate has improved to 70%, the combined treatment can cause severe side effects and late complications in survivors, such as cognitive impairment, endocrine disorders, and increased risk of secondary tumors, especially in young children [1]. To reduce treatment-related side effects while maintaining treatment efficacy, efforts have been made to find a targeted therapy for SHH-MB [16]. Vismodegib (GDC-0449) and Sonidegib (LDE225) were developed as SMO inhibitors and approved for treatment of advanced basal cell carcinoma (BCC) and entered clinical trials for SHH-MB in combination with conventional chemotherapies [17, 18]. Studies have shown that Vismodegib and Sonidegib are effective for relapsed SHH medulloblastoma in adults [19, 20]. However, in infants and young children, inhibition of SMO can lead to toxicity in multiple organs, particularly inhibition of bone growth, which has been shown to be irreversible in mice [21–23]. In addition, the long-term efficacy of SMO inhibition is limited due to the emergence of drug resistance, and patients with mutations downstream of SMO may not benefit [11, 24]. Therapies targeting other signaling pathways are still under investigation [16]. Overall, this underscores the urgent need for improved targeted therapies that minimize harmful side effects for the treatment of MB. To find more effective treatments for SHH-MB while sparing normal neuronal cells, we developed a high-throughput screening platform using our previously established neuroepithelial stem (NES) cell model [25] and assayed a library of compounds of known targets identifying S6K1 as a selective target for SHH-MB.

## Materials and Methods

### Cell culture

Generation of Control NES, Parental NES and 2^nd^ tNES cells was previously described in [25]. NES cells were grown in DMEM/F12 media supplemented with B27, N2, EGF and FGF2. DAOY, UW2283 and HEK293FT cells were grown in DMEM media with 10% FBS. ONS76 cells were grown in RPMI media with 10% FBS. See Supplementary Methods for detailed information.

### DiSCoVER analysis method

The DiSCoVER method is publicly available as an analysis module in GenePattern (https://www.genepattern.org). DiSCoVER predicts drug sensitivity through comparing gene expression profile of target cell and cancer cell lines with known drug sensitivity data. See Supplementary Methods for detailed information.

### High-throughput compound screening platform

We developed a high-throughput cytotoxicity plate based-assay using the Cell Titer-Glo® 2.0 Luminescent Cell Viability Assay (Promega, G9242) to screen CBCS Oncoset collection of 172 compounds with known targets (listed in table S1) to identify compounds that selectively inhibit the growth of 2^nd^ tNES with the Parental NES as the control reference. Parental NES (2^nd^ tNES) were plated at 8000 (4000) cells per well in 384-well plate (PerkinElmer, 6007480) pre-coated with 0.1 mg/ml poly-L-ornithine (Sigma, P3655-100mg). Laminin (Sigma, L2020) was added to each well to achieve the final concentration of 1 μg/ml. For the primary screen, compounds were tested at 1μM and 10μM. Then the hits (compounds inhibit cell viability of 2^nd^ tNES by more than 20%) were retested in full dose-response (1:2 serial dilution from 20nM - 20μM) and selectivity were subsequently confirmed (1:3 serial dilution from 0.2nM - 20μM). After 48 hours incubation with the compounds, cell viability was measured using the Cell Titer-Glo® 2.0 Luminescent Cell Viability Assay. Nonlinear regression (four-parameter variable point equation) was performed for EC50 calculation (GraphPad Prism 9.0). DMSO (0.1%) was used as negative control (100% viability) and Etoposide (10μM) was used as positive control (0% viability). For each plate, Z factor, signal to background ratio, and coefficient of variation were monitored to evaluate assay performance and data quality.

### Single agent and combination treatment

Parental NES, 2^nd^ tNES, Daoy, UW2283 and ONS76 were seeded in 96-well plate and incubated with Brivanib and PF4708671 either alone or combined with Cyclophosphamide or Vincristine for 48 hours. Resazurin sodium salt (Sigma, R7017) was used to assess cell viability. Cell seeding density and compound concentration varies between cell lines (Table S4, S5). See Supplementary Methods for detailed information.

### Cytotoxicity assay

Parental NES and 2^nd^ tNES were seeded in 96-well plate and incubated with Brivanib and PF4708671 for 48 hours. Cytotoxicity was assessed CytoTox 96® Non-Radioactive Cytotoxicity Assay kit (Promega, G1780) according to manufacturer’s instructions. See Supplementary Methods for detailed information.

### Immune cell proliferation

CD8+ T cells and γδ T cells were isolated from PBMC from healthy donors and incubated with Brivanib or PF4708671 for 72 hours. Cell proliferation was measured by flow cytometry. See Supplementary Methods for detailed information.

### Western blot

Western blot was performed as previously described [25]. See Supplementary Methods for detailed information.

### Immunofluorescence staining

Neurons differentiated from Parental NES were stained for Beta III tubulin and cleaved caspase 3. Samples were analyzed under the fluorescent microscope Zeiss LSM900. Staining intensities were quantified by ImageJ software. See Supplementary Methods for detailed information.

### Cell cycle analysis

Cells were stained with propidium iodide and analyzed by flow cytometry. Cell cycle distribution analysis was performed using FlowJo (FlowJo LLC, Ashland, OR). See Supplementary Methods for detailed information.

### 3D Spheroid assay

Parental NES and 2^nd^ tNES were seeded at 3000 cells per well in ultra-low attachment U-bottom 96 well plate (FisherScientific, 15227905) and incubated with Brivanib (360nM) and PF4708671 (750nM and 1.4μM) for 3 days. The size of spheroids was analyzed to reflect the prophylactic effect of treatment. To assess the therapeutic effect, spheroids were formed in normal culture medium for 3 days followed by treatment at same concentrations for another 3 days. Image analysis was performed with INSIDIA macro in ImageJ [26].

### Lentiviral shRNA-mediated knockdown of RPS6KB1

Lentiviral particles were made by co-transfection of HEK293FT cells with expression vectors, packaging plasmid psPAX2 and pMD2G in a ratio of 2:2:1 using lipofectamine 3000 (ThermoFisher, L3000001) and transduced into NES cells and ONS76 cells. Selection was performed using puromycin and knockdown efficiency was assessed by protein level. See Supplementary Methods for detailed information.

### Zebrafish experiment

*In vivo* treatment of PF4798671 was performed in zebrafish as described in van Bree et al. in prep. See Supplementary Methods for detailed information.

### Mice experiment

ONS76 cells and 2^nd^ tNES were knocked down of RPS6KB1 gene and subcutaneously (2×10^6^ ONS76 cells) and orthotopically (50,000 2^nd^ tNES) injected into NSG mice. Tumor growth was captured with calipers or bioluminescence measurements. See Supplementary Methods for detailed information.

### Statistical analyses

Unpaired two-tailed Student’s t tests were used to evaluate the statistical significance of differences between treatment groups for flow cytometry data, spheroid data and LDH data, where a P value <0.05 was considered statistically significant. All data analysis was performed using GraphPad Prism 9. Values of significance are indicated by asterisks (* p<0.05, ** p<0.01, *** p<0.001, **** p<0.0001) and described in each Fig. legend where appropriate.

## Results

### Combined analysis of gene expression patterns and drug sensitivity profiles identify multiple signaling pathways as potential targets for SHH-MB treatment

We have previously developed a human SHH-MB stem cell model by reprogramming non-cancerous cells from a Gorlin syndrome patient carrying a germline *PTCH1* mutation (1762insG) into induced pluripotent stem (iPS) cells and differentiating them into neuroepithelial stem (NES) cells [25]. We have shown that patient-derived NES cells (termed Parental NES) form MB tumors when orthotopically injected into the cerebellum of NSG mice, with malignancy increasing with each round of injection (Fig. 1A) [25]. Differential gene expression profiles of Parental NES and tumor NES isolated from tumors after two rounds of cerebellar injection (termed 2^nd^ tNES) may provide insight into targetable pathways that drive malignant conversion of Parental NES to 2^nd^ tNES *in vivo.* Therefore, we compared their gene expression profiles and used the DiSCoVER algorithm [27] to predict drug sensitivity. Briefly, DiSCoVER generates a 2^nd^ tNES oncogenic enrichment signature (OES) by comparing gene expression profile of 2^nd^ tNES with Parental NES and predicted drug sensitivity by comparing the 2^nd^ tNES signature with gene expression profiles of different cancer cell lines with known drug response data (Fig. 1B). The resulting drug sensitivity predictions were quantified using a score, where higher scores indicate likely sensitivity, and lower or negative scores indicate potential resistance (details in Supplementary Materials and Methods). Moreover, to compare the predicted drug sensitivity between 2^nd^ tNES and patient SHH-MB, we generated a patient SHH-MB OES by re-analyzing a microarray expression dataset of 405 SHH-MB and 291 normal cerebellum/upper rhombic lip (uRL) [28]. The 2^nd^ tNES OES and patient SHH-MB OES were then compared with two different drug sensitivity datasets, GDSC (Genomics of Drug Sensitivity in Cancer) and CTRPv2 (The Cancer Therapeutics Response Portal), to suggest potentially effective drugs against 2^nd^ tNES and patient SHH-MB. The predicted drugs were clustered based on mechanism-of-action (MoA) and within each category ranked by a score which is positively correlated with drug efficiency (Figs. 1C-E, fig. S1). From the GDSC dataset, about 50% of compounds targeting cell cycle/DNA damage, epigenetic modifiers, MAPK/ERK, PI3K/AKT/mTOR and receptor tyrosine kinase (RTK) were predicted to be effective against 2^nd^ tNES cell. Most of them were also predicted to be effective against patient SHH-MB (Figs. 1C-E, figs. S1A-B). From the CTRPv2 dataset, approximately 50% of compounds targeting MAPK/ERK, PI3K/AKT/mTOR predicted efficacy against both 2^nd^ tNES cells and patient SHH-MB (Figs. 1D-E, figs. S1C-D). Taken together, these results suggest that the NES model mimics SHH-MB patients in predicting potential therapeutics. Moreover, it confirms previous studies on the importance of targeting commonly dysregulated signaling pathways in MB such as MAPK/ERK, PI3K/AKT/mTOR, and RTK signaling pathways as potential treatment options for SHH-MB [29–31].

**Figure 1.**
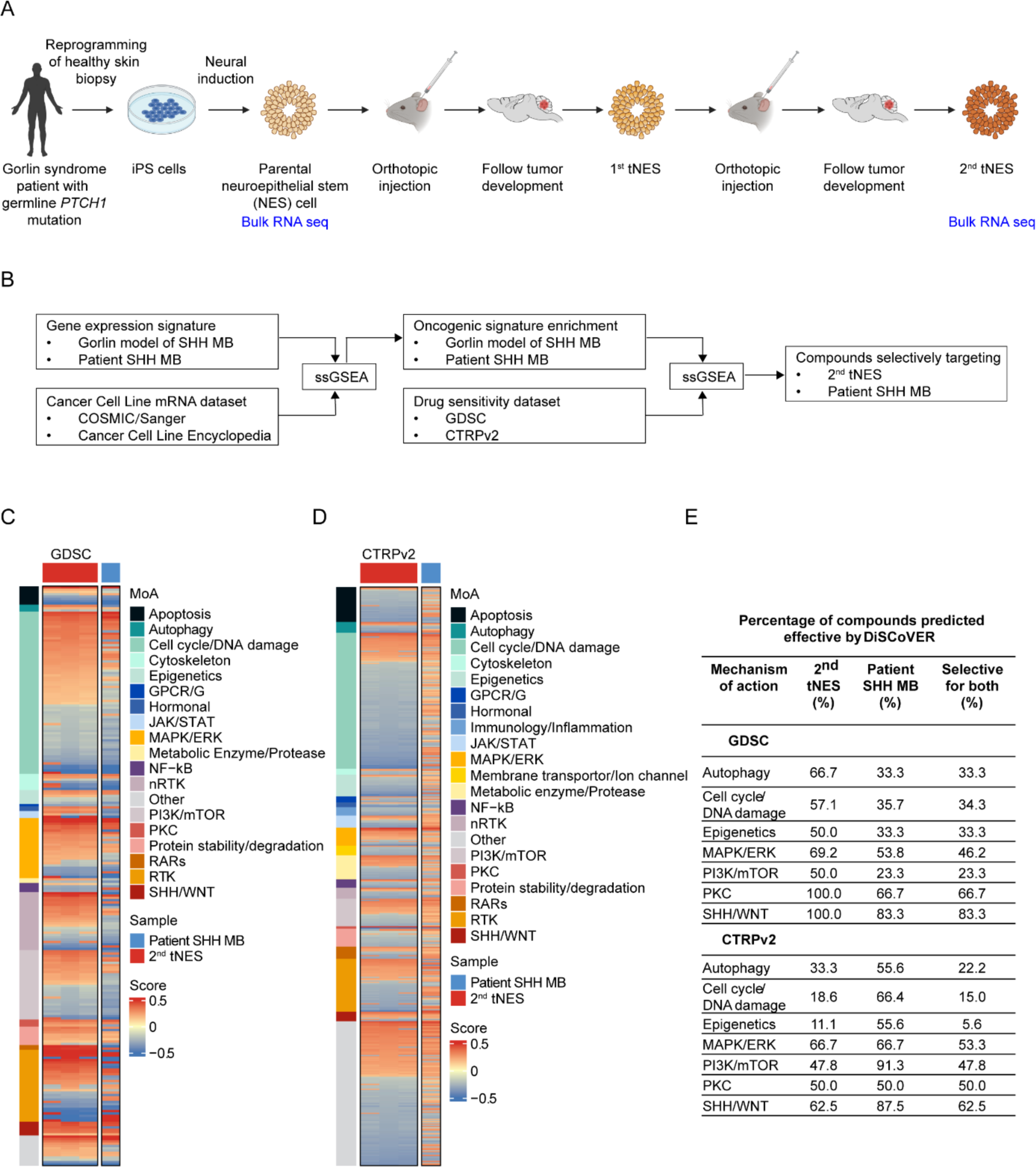
DiSCoVER algorithm predicts potential targets of SHH medulloblastoma treatment. (A) Schematic overview of neuroepithelial stem (NES) cell model establishment and tNES cell generation through isolation and orthotopic reinjection. (B) Diagram showing main steps in the DiSCoVER analysis method. (C) Compound predicted from GDSC dataset clustered by MoA and ranked by efficacy against 2^nd^ tNES. (D) Compound from CTRPv2 dataset clustered by MoA and ranked by efficacy against 2^nd^ tNES. (E) Summary table showing individual and overlapped percentage of compounds predicted efficient against 2^nd^ tNES from GDSC and CTRPv2 datasets.

### High-throughput compound screening identifies SHH-MB sensitivity to compounds targeting p70S6K1 and VEGFR2/FGFR1

To validate the biological pathways predicted by DiSCoVER and identify new potential therapeutics, we developed a plate-based high-throughput viability assay and screened the 2^nd^ tNES (#1440), Parental NES, and NES cells derived from a healthy individual (Ctrl1) with an Oncoset library consisting of 172 known compounds targeting receptor or non-receptor RTKs, PI3K/mTOR, MAPK/ERK, hormone receptors, DNA damage, cell cycle and other pathways (Fig. 2C, Table S1). Compounds efficacy and selectivity were determined based on cell viability and differences between Parental NES and 2^nd^ tNES, which share the same genetic background (Figs. 2A-B). Compounds were first screened at 1μM and 10μM and DMSO and etoposide were used as negative and positive controls, respectively.

**Figure 2.**
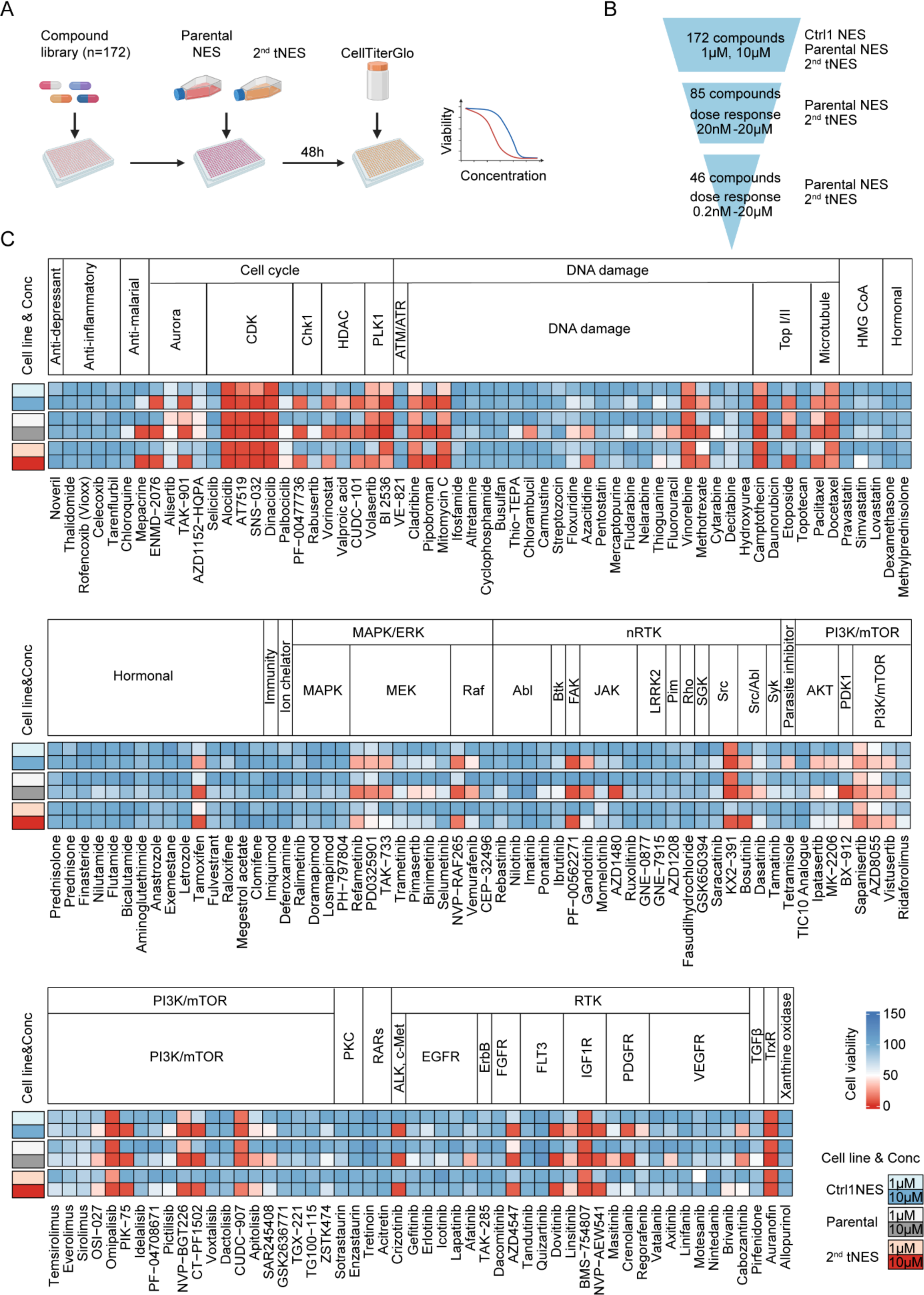
High-throughput compound screening identifies potential compounds for SHH-MB treatment. (A) Schematic overview of screening setup. (B) Funnel diagram of screening process. (C) Heatmap showing cell viability of Ctrl NES, Parental NES and 2^nd^ tNES treated with a library of 172 compounds at 1μM and 10μM.

Of the 172 compounds, 85 showed efficacy on 2^nd^ tNES, including inhibitors targeting CDK and PLK1 (Cell cycle pathway), MEK and RAF (MAPK/ERK pathway), PI3K/mTOR (PI3K/mTOR pathway), FGFR, IGF1R and VEGFR (RTK pathway) and DNA damaging pathway. However, most of these compounds showed similar efficacy on Parental NES and Ctrl1 NES cells, suggesting that these pathways are important for the survival of both tumor cells and normal neural stem cells (Fig. 2C). As these compounds were tested at only two concentrations and many of them appeared extremely toxic, to further evaluate the selectivity of the compounds between Parental NES and 2^nd^ tNES, 85 compounds that inhibited the cell viability of the 2^nd^ tNES by more than 20% were included for a 11-point dose-response test (20nM to 20μM in singlet) (Fig. S2, Table S2). For each compound a dose-response curve was determined based on cell viability and a drug sensitivity score (DSS) was calculated based on the area under the dose-response curve (AUC). Selectivity was determined by the differential cell viability and DSS (dDSS) between Parental NES and 2^nd^ tNES [32]. Compounds with dDSS above 0 indicates selectivity for 2^nd^ tNES, dDSS equal to 0 indicates no selectivity and dDSS below 0 indicates selectivity for Parental NES. The top ranked compounds selective for 2^nd^ tNES were Decitabine (DNA damage), Azacitidine (DNA damage), Brivanib (VEGFR/FGFR), Alisertib (Cell cycle), Masitinib (PDGFR) and PF4708671 (p70S6K1) (Fig. 3A).

**Figure 3.**
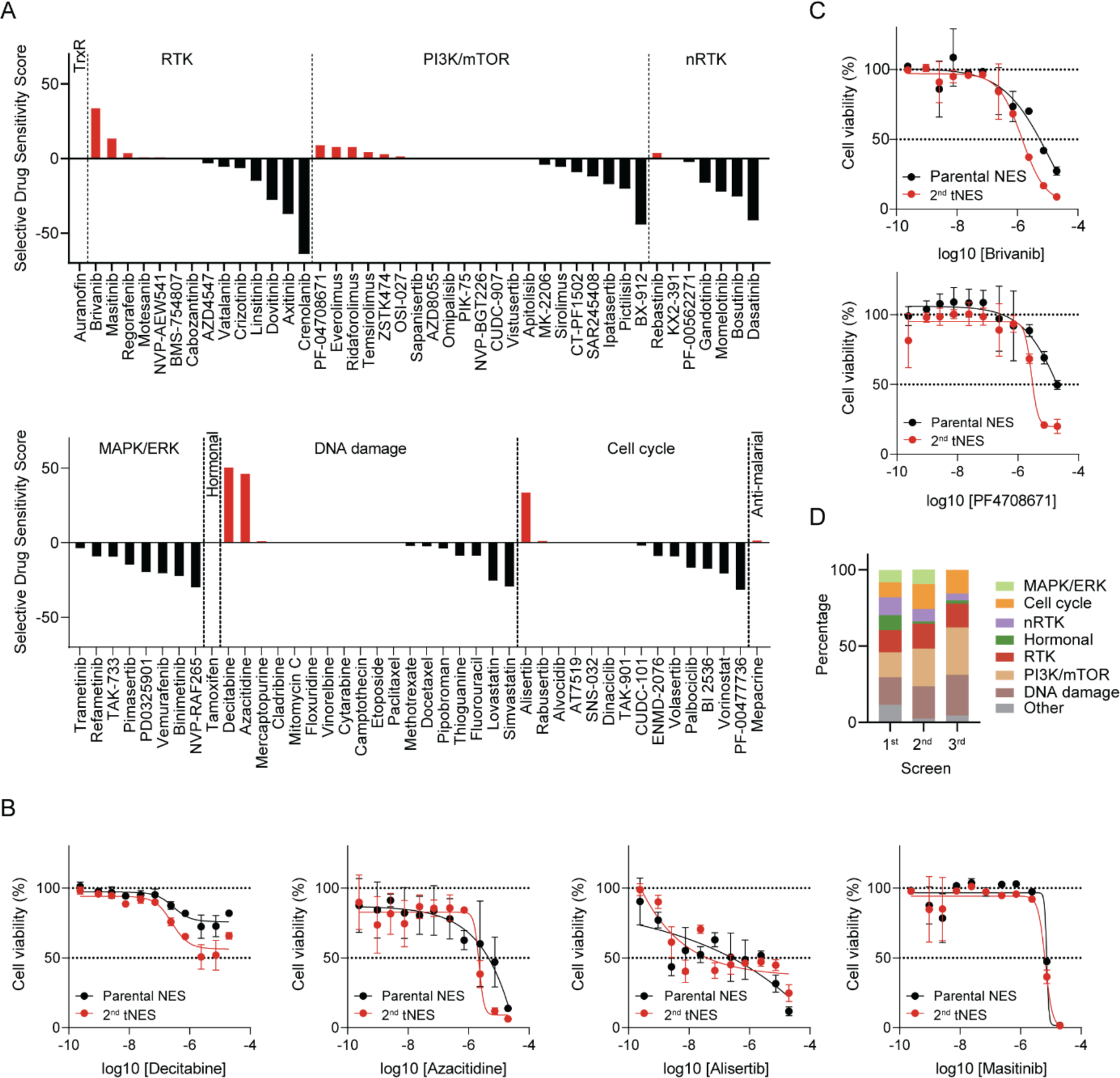
Dose-response validation identifies compound selectivity between Parental NES and 2^nd^ tNES (A) Selective Drug Sensitivity Score (sDSS) of 85 compounds between Parental NES and 2^nd^ tNES. sDSS > 0 indicates selectivity toward 2^nd^ tNES, sDSS = 0 indicates no selectivity, sDSS < 0 indicates selectivity toward Parental NES. (B) Cytotoxic dose response curve of Decitabine (EC50 _Parental NES_ = 0.26μM, EC50 _2_^nd^ _tNES_ = 0.29μM), Azacitidine (EC50 _Parental NES_ = 2.69μM, EC50 _2_^nd^ _tNES_ = 1.75μM), Alisertib (EC50 _Parental NES_ = 0.13μM, EC50 _2_^nd^ _tNES_ = 0.22μM), and Masitinib (EC50 _Parental NES_ = 7.16μM, EC50 _2_^nd^ _tNES_ = 5.97μM) derived from 11-point dose-response screening in triplicate. (C) Cytotoxic dose response curve of Brivanib and PF4708671 derived from 11-point dose-response screening in triplicate. (D) Composition of compound library of primary screening (1^st^) and dose-response in singlet (2^nd^) and in triplicate (3^rd^)

Based on the difference in dDSS between Parental NES and 2^nd^ tNES, we included 46 compounds that showed selectivity towards 2^nd^ tNES for confirmation in 11-point dose-response manner (0.2nM to 20μM in triplicate) (Fig. S3, Table S3). The EC50 of compounds in Parental NES and 2^nd^ tNES were analyzed to identify selective hit compounds. Although Decitabine, Azacitidine, Alisertib and Masitinib had favorable dDSS, they showed poor efficacy in drug response or a narrow therapeutic window (Fig. 3B). However, combined analysis identified Brivanib (VEGFR1/FGFR2 inhibitor) and PF4708671 (p70S6K1 inhibitor) as selective towards 2^nd^ tNES with a EC50 compared to Parental NES 8.6-fold and 13.4-fold higher, respectively (Fig. 3C).

VEGF receptors and FGF receptors have been found to be expressed on MB cell lines and PDX cells, and inhibition of VEGFR and FGFR can suppress tumor cell growth and invasive potential [33–35]. S6K1 is a downstream target of PI3K/mTOR pathway and interestingly, by analyzing the composition of the library in the screening, we found that compounds targeting the PI3K/mTOR pathway were significantly enriched, which indicated the potential vulnerability of 2^nd^ tNES to PI3K/mTOR inhibitors (Fig. 3D). Previous studies have shown that the PI3K/mTOR pathway is frequently activated in MB and its crosstalk with other pathways may promote tumorigenesis [31, 36, 37]. Compounds developed to inhibit key components of PI3K/mTOR pathway have shown promising preclinical antitumor effects and have been enrolled in clinical trials [38]. Although the role of S6K1 in MB remains to be determined, S6K1 was found to be associated with tumor growth and poor prognosis in different types of cancer [39–42] and S6K1 inhibition attenuated drug resistance to palbociclib [43] and EGFR TKIs [40]. However, as a key component of the PI3K/mTOR pathway, S6K1 has been studied less in the pathogenesis and therapy of MB. Taken together, the *in vitro* screening suggests that compounds targeting S6K1 and VEGFR1/FGFR2 would act specifically on SHH-MB tumor cells and not on normal neural stem cells.

### Brivanib and PF4708671 inhibit the growth of MB tumor cells and act synergistically in combination with conventional chemotherapy

To validate the selectivity of Brivanib and PF4708671, we included three control NES cell lines derived from healthy individuals (Ctrl1, Ctrl7 and Ctrl9) [44–46], Parental NES and three 2^nd^ tNES (#1440, #1463 and #1471) for cell viability analysis. Based on the dose-response curve and EC50 value, all control NES cells showed similar sensitivity to Brivanib and PF4708671 as Parental NES whereas all 2^nd^ tNES cells showed greater sensitivity to Brivanib and PF4708671 than both Ctrl NES and Parental NES (Fig. 4A).

**Figure 4.**
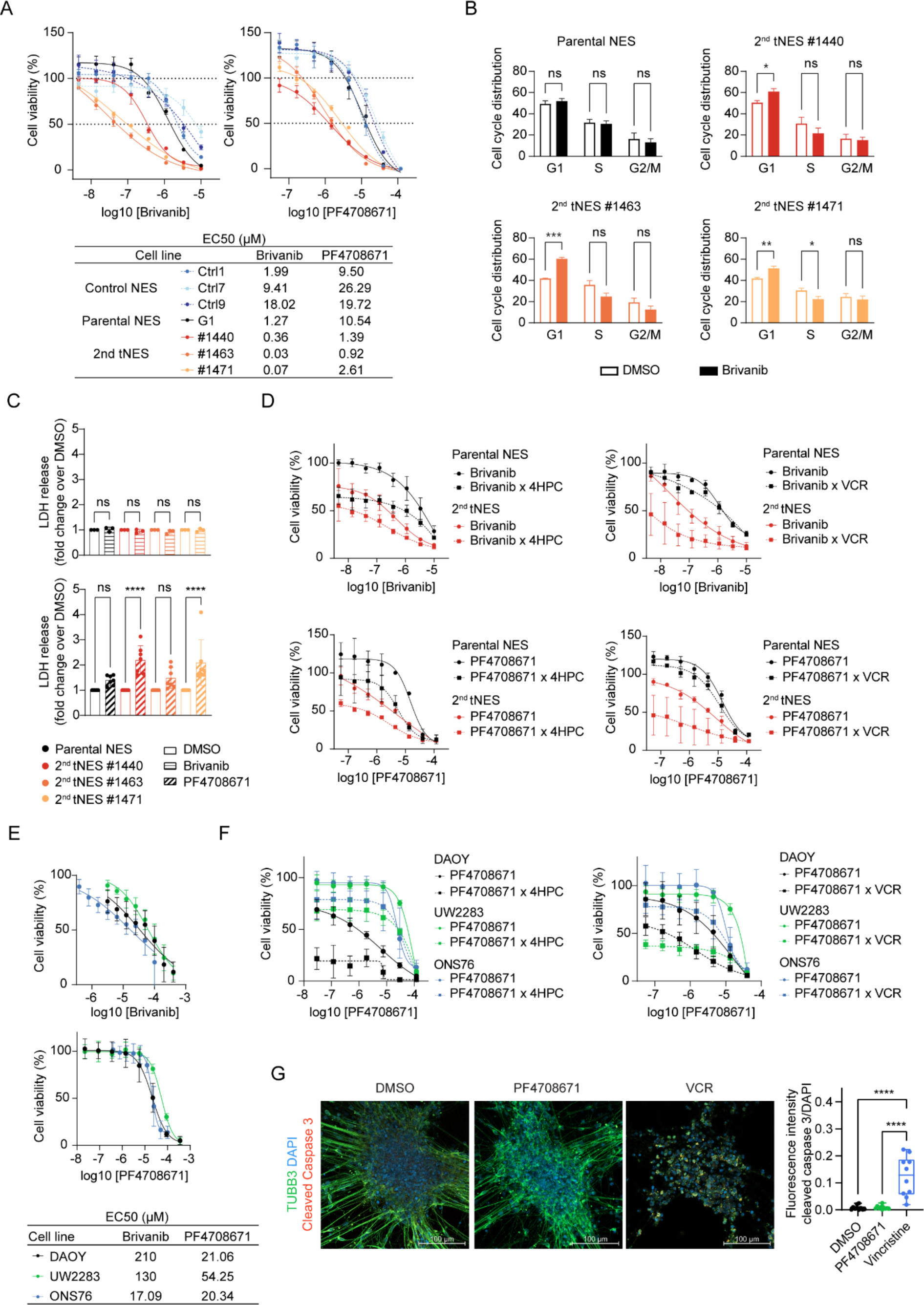
Brivanib and PF4708671 show selectivity towards 2^nd^ tNES in functional assays. (A) Cell viability of Ctrl NES (Ctrl1, Ctrl7, Ctrl9), Parental NES and 2^nd^ tNES (#1440. #1463, #1471) treated with Brivanib and PF4708671 (measured with resazurin, n=3 independent experiments). (B) Cell cycle analysis of Parental NES and 2^nd^ tNES treated with Brivanib (n=4 independent experiments). (C) Cytotoxicity assay measured by LDH level in supernatant of Parental NES and 2^nd^ tNES treated with Brivanib and PF4708671 (n=3-7 independent experiments) (D) Cell viability of Parental NES and 2^nd^ tNES treated with combination of Brivanib or PF4708671 and 4HPC (0.2μM) or VCR (4.4nM) (n=3 independent experiments). (E) Cell viability of Daoy, UW2283 and ONS76 cells treated with Brivanib and PF4708671 (n=3 independent experiments). (F) Cell viability of Daoy, UW2283 and ONS76 cells treated with combination of PF4708671 and 4HPC (12.5μM) or VCR (7.5nM) (n=3 independent experiments). (G) Immunofluorescent staining of beta tubulin III and cleaved caspase 3 in Parental NES differentiated neurons treated with DMSO, PF4708671 (1.4μM) and Vincristine (40nM) and quantification of cleaved caspase 3. Each dot represents one image in the same experiment.

Next, we analyzed whether the decrease in cell viability was due to decreased proliferation or increased cell death. Cell cycle analysis showed that Brivanib caused significant G1 cell cycle arrest only in 2^nd^ tNES, whereas PF4708671 had no effect on the cell cycle in either Parental NES or 2^nd^ tNES (Fig. 4B, fig. S9A). In contrast, a lactate dehydrogenase (LDH) release assay measuring cell death showed no increase in LDH release with Brivanib in the parental NES or 2^nd^ tNES, whereas a significant increase in LDH release was detected with PF478671 in the 2^nd^ tNES compared with Parental NES (Fig. 4C), suggesting that PF4708671 has cytotoxic activity, whereas Brivanib is mainly cytostatic.

Considering that cyclophosphamide and vincristine (VCR) have been commonly used in chemotherapy for MB, we next examined the effect of combination treatment in Parental NES and 2^nd^ tNES to assess the interaction between Brivanib or PF4708671 and cyclophosphamide or VCR. Cyclophosphamide is a prodrug; therefore, we used the activated form 4-hydroperoxycyclophospamide (4HPC). SynergyFinder was used to evaluate combinatorial activity of drugs, with a synergy score of less than - 10 considered antagonistic, from −10 to 10 considered additive, and greater than 10 considered synergistic [47].

Importantly, 2^nd^ tNES were more sensitive than Parental NES to all combination treatments of Brivanib or PF4708671 together with 4HPC or VCR, as reflected by cell viability (Fig. 4D, Fig. S4, S5). Brivanib and 4HPC acted mainly additive and synergistically at only a few dose combinations in the 2^nd^ tNES compared with the Parental NES (Fig. S4A), whereas the combination of Brivanib and VCR showed more synergy than the Parental NES at relatively low concentrations in the 2^nd^ tNES (Fig. S4B). Combinations of PF408671 and 4HPC or VCR showed synergy at most concentration combinations in the 2^nd^ tNES, which was observed in the Parental NES only at very high concentrations of PF4708671 or VCR (Fig. S5). These results showed that Brivanib and VCR generally acted synergistically in the 2^nd^ tNES, and PF4708671 acted synergistically with both 4HPC and VCR in 2^nd^ tNES, whereas it acted less synergistically or antagonistically in the Parental NES.

We next tested the efficacy of Brivanib or PF4708671 in established SHH-MB cell lines as monotherapy or combination therapy with 4HPC and VCR. DAOY, UW2283, and ONS76 cells were sensitive to both Brivanib and PF4708671 as monotherapy, although not to the same extent as tNES cells (Fig. 4E). However, we observed a significant synergistic effect with PF4708671 as combination treatment with 4HPC or VCR at certain concentrations (Fig. 4F, Fig. S6). In addition, combination treatment with Brivanib and PF4708671 acted antagonistically in the Parental NES but synergistically in the 2^nd^ tNES, DAOY, and UW2283 cells (Fig. S7). Taken together, these results suggest that Brivanib and PF4708671 selectively inhibit MB tumor cell growth and have a minor effect on normal neural stem cells. PF4708671 acted synergistically with cyclophosphamide and VCR, and the combination treatment showed selectivity toward tumor cells.

### Brivanib and PF4708671 do not induce treatment-related neurotoxicity or immunotoxicity

Both conventional and novel cancer therapies can cause treatment-related neurotoxicity resulting in patient morbidity and mortality. To evaluate the safety profile on neurons, we evaluated the expression of cleaved caspase 3 and the neurite structure of neurons differentiated from Parental NES cells. Whereas VCR, that was used as the positive control, damaged neurite structure and significantly induced cleaved caspase 3 expression compared to DMSO, treatment with either Brivanib or PF470867 at their EC50 concentration in 2^nd^ tNES had no visible effects neurons (Fig. 4G, fig. S8B).

While targeted therapy shows an anti-tumor effect on tumor cells, it could modulate the immune system to provide an additional anti-tumor effect or counteract the effect of targeted therapy [48]. To test the effect on immune response, we treated human CD8+ T cells and γδT cells, which are the main T cell effectors in cancer immunotherapy, with Brivanib and PF4708671 and found no effect on cell survival or proliferation (Figs. S8C, S8D). Altogether showing that Brivanib and PF4708671 do not induce neurotoxicity or affect immune effector cell proliferation.

### PF4708671 impairs MB tumor cell growth in 3D culture and in a zebrafish MB model

Cancer cells cultured in two-dimensional (2D) environments tend to proliferate more rapidly, exhibit different gene and protein expression and metabolic profiles, and lack cell-cell and cell-matrix interactions, which could alter drug sensitivity. To overcome these limitations, three-dimensional (3D) cancer models have been developed, and the tumor spheroid model has been widely used to study drug response and efficacy [49, 50]. We first tested the effects on 3D formation by allowing spheroid formation in the presence of Brivanib or PF4708671. Surprisingly, Brivanib had no effect on spheroid formation of Parental NES or 2^nd^ tNES (Fig. 5A). In contrast, PF4708671 significantly decreased the size of spheroids of 2^nd^ tNES at low (750μM) and high (1.4uμM) concentrations, while only the high concentration affected spheroids of Parental NES (Figs. 5B-C). Next, we tested the efficacy on already established spheroids. Again, Brivanib had no measurable effect on tumor spheroid size (Fig. S8E), whereas PF4708671 significantly reduced 2^nd^ tNES spheroid size, while Parental NES spheroid size was unaffected (Fig. S8E).

**Figure 5.**
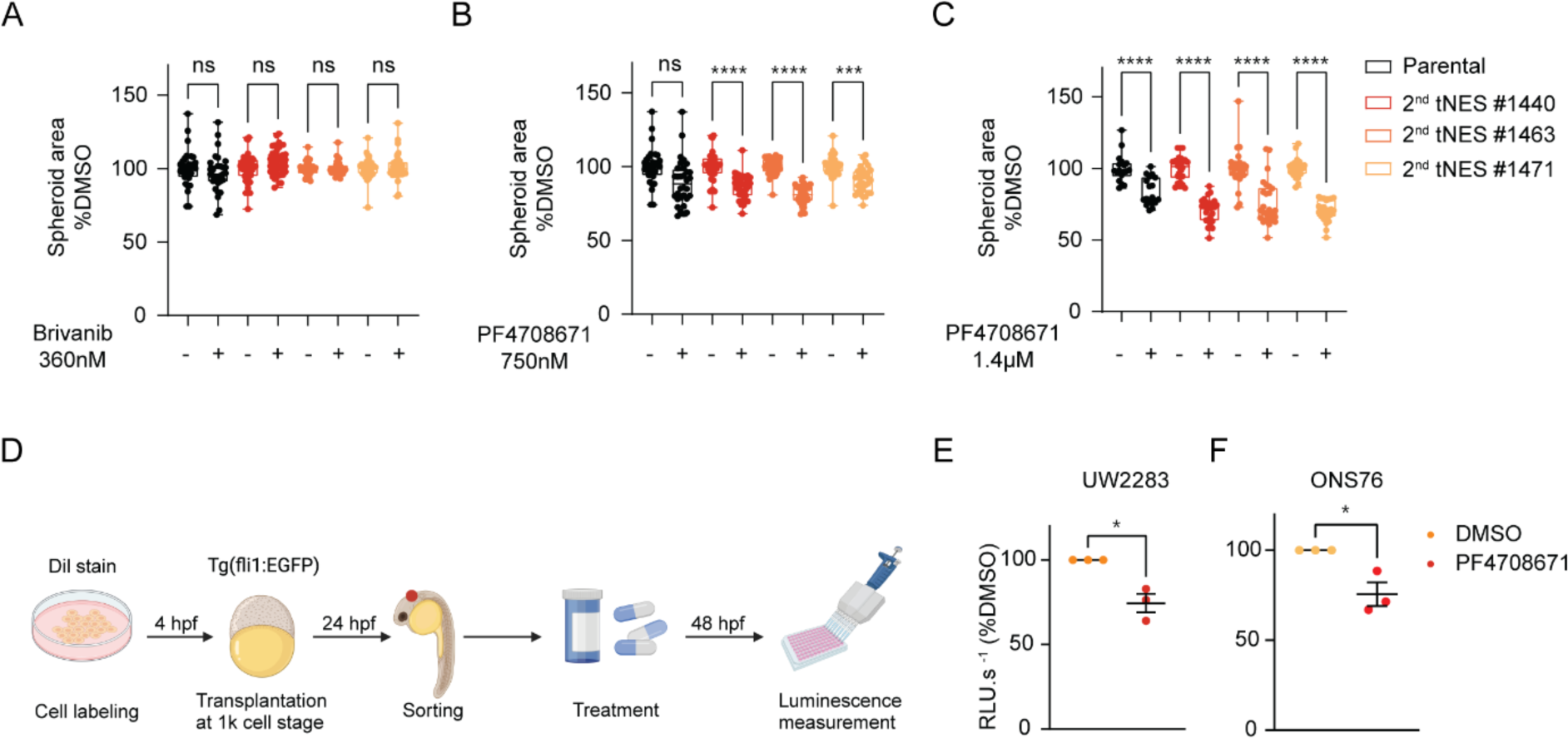
Effect of PF4708671 in tumor spheroid formation and zebrafish xenograft model. (A) Boxplot of size of spheroid derived from Parental NES and 2^nd^ tNES upon Brivanib (360nM) treatment (n=4). (B) Boxplot of size of spheroid derived from Parental NES and 2^nd^ tNES upon PF4708671 (750nM) treatment (n=4). (C) Boxplot of size of spheroid derived from Parental NES and 2^nd^ tNES upon PF4708671 (1.4μM) treatment (n=3). (D) Schematic overview of PF4708671 treatment in zebrafish xenograft model. (E) *In vivo* cell viability of UW228-3-Luc upon 48h PF4708671 treatment (10μM) in zebrafish xenograft model (each dot represents mean value from each experiment, n=3 independent experiments, in total 52 DMSO and 54 PF4708671 zebrafish embryos were analyzed).(F) *In vivo* cell viability of ONS76-Luc upon 48 h PF4708671 treatment (10μM) in zebrafish xenograft model (each dot represents mean value from each experiment, n=3 independent experiments, in total 36 DMSO and 35 PF4708671 zebrafish embryos were analyzed).

Since Brivanib showed only cytostatic activity and had no measurable effect on cells growing in a 3D environment, we focused our study on targeting S6K1. To further evaluate the anti-tumor effect of S6K1 inhibition, we used a recently established MB zebrafish embryo model in which medulloblastoma cells are transplanted into zebrafish blastomeres at the 1k cell stage and migrate to form a cell mass in the hindbrain region of the developing nervous system (van Bree et al. in prep). Dil-stained UW228-3-Luc and ONS76-Luc cells were transplanted, drug treatment was started 24 hours after transplantation, and bioluminescence activity was measured to reflect the number of living tumor cells. The result showed that PF4708671 significantly decreased the cell viability of UW228-3 and ONS76 cells by 25% and 24% respectively. (Fig. 5D-F). Taken together, these results suggest that PF4708671 inhibits tumor spheroid formation *in vitro* and tumor cell growth *in vivo*.

### Increased PI3K/AKT signaling sensitizes medulloblastoma cells to S6K1 inhibition

To identify the mechanism underlying the selectivity of PF4708671, we analyzed the gene expression profile and found that the PI3K/AKT pathway was enriched in 2^nd^ tNES compared with Parental NES (Fig. 6A). The upregulation of the PI3K/AKT pathway was also found in SHH-MB patients (Fig. 6B), as previously reported [51, 52]. To test the correlation between the PI3K/AKT pathway and sensitivity to PF4708671, we divided 268 cancer lines from the GDSC database into PF4708671-sensitive (EC50 below 60μM) and PF4708671-resistant (EC50 above 60μM) cells to perform differential gene expression and pathway analysis, and we found that the PI3K/AKT pathway was enriched in the PF4708671-sensitive cells (Fig. 6C). In addition, gene set enrichment analysis of gene expression profiles comparing Parental NES with 2^nd^ tNES revealed enrichment of the mTORC1 pathway in the 2^nd^ tNES (Fig. 6D). Taken together, these results suggest that the PI3K/AKT pathway is upregulated in SHH-MB and correlates with sensitivity to PF4708671.

**Figure 6.**
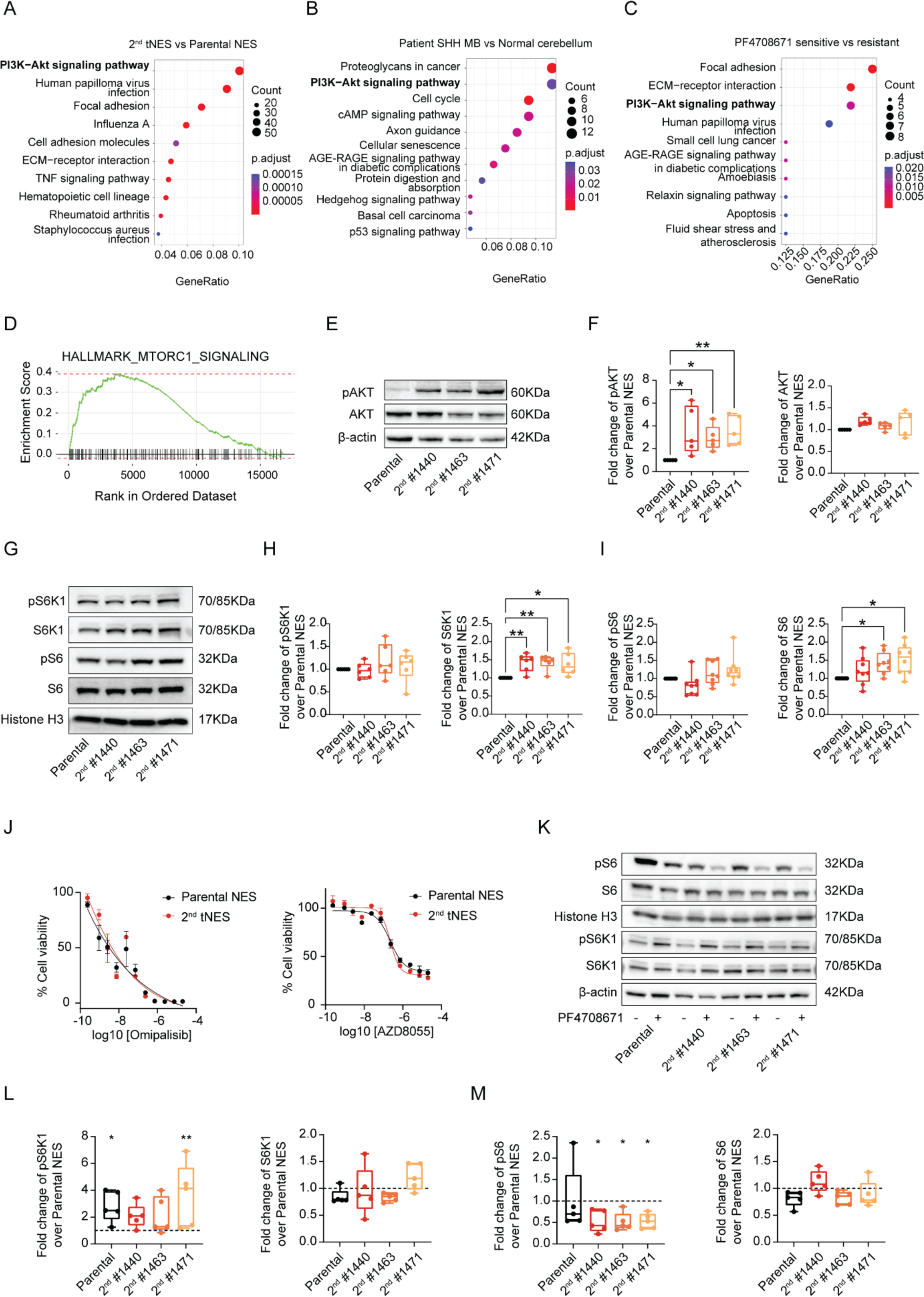
PI3K/AKT signaling pathway activation in 2^nd^ tNES and S6K1 signaling inhibition upon PF4708671 treatment. (A) Pathways enriched in 2^nd^ tNES compared to Parental NES by KEGG analysis. (B) Pathways enriched in SHH-MB patients compared to normal cerebellum by KEGG analysis. (C) Pathways enriched in PF4708671 sensitive (EC50 <= 60uM, n=166) cell lines compared to PF4708671 resistant (EC50 > 60uM, n=66) cell lines by KEGG analysis. (D) Hallmark mTORC1 signaling enriched in 2^nd^ tNES compared to Parental NES by GSEA analysis. (E) Representative Western blots of protein level of total and phosphorylated S6K1, S6, AKT and ERK in Parental NES and 2^nd^ tNES. (F-I) Quantification of protein level of total and phosphorylated S6K1, S6, AKT and ERK in Parental NES and 2^nd^ tNES (n=5-8). (J) Dose-response curve of Omipalisib and AZD8055 in Parental NES and 2^nd^ tNES from dose-response screening in triplicate. (K) Representative Western blots of protein level of total and phosphorylated S6K1 and S6 in Parental NES and 2^nd^ tNES upon DMSO or PF4708671 treatment. (L-M) Quantification of protein level of total and phosphorylated S6K1 and S6 in Parental NES and 2^nd^ tNES (n=5).

We then confirmed the enhanced PI3K/AKT activity in 2^nd^ tNES by measuring total and phosphorylated AKT protein levels and found significant upregulation of phospho-AKT, total S6K1 and total S6 protein in 2^nd^ tNES compared with Parental NES (Figs. 6E-I). Surprisingly, we did not detect significant differences in phosphorylated S6K1 or S6 between Parental NES and 2^nd^ tNES (Figs. 6G-I). Moreover, blocking PI3K, mTOR or both did not selectively reduce cell viability of 2^nd^ tNES compared with Parental NES (Fig. 6J, fig. S9). After PF4708671 treatment, we observed increased expression of phospho-S6K1 in both Parental NES and 2^nd^ tNES, consistent with previous observations [53]. However, PF4708671 treatment resulted in a greater decrease in phospho-S6 in the 2^nd^ tNES compared to Parental NES (Figs. 6K-M). Taken together, these results suggest increased PI3K/AKT pathway activation in the 2^nd^ tNES compared with Parental NES, which is consistent with previous studies showing activation of the PI3K/AKT pathway in MB [51]. And blocking S6K1, a downstream target in the PI3K/AKT pathway, resulted in selectivity against SHH-MB tumor cells compared to normal neural stem cells.

### S6K1 knockdown impairs SHH MB tumor cell viability and *in vivo* tumor growth

To confirm the selective susceptibility of tumor cells to S6K1 inhibition, we tested the effect of small hairpin RNA (shRNA)-mediated S6K1 (*RPS6KB1*) knockdown (KD) on cell viability in Parental NES and 2^nd^ tNES using two different shRNAs targeting S6K1. The knockdown efficacy was assessed by S6K1 protein level. Through measuring the cell viability, we found that 2^nd^ tNES was more sensitive to S6K1 KD than Parental NES (Figs. 7A-B). The effect of S6K1 KD on cell proliferation could be validated in the ONS76 cell line (Figs. 7C-D), suggesting that tumor cells were more affected by S6K1 KD than normal neural stem cells. To test the effect of S6K1 KD *in vivo*, ONS76 shCtrl and shS6K1 cells were subcutaneously injected in NSG mice and followed for tumor development. Knock down of S6K1 significantly impaired tumor growth over time (Fig. 7E) and tumor size (Fig. 7F). The effect of S6K1 KD on tumor growth was verified by orthotopic cerebellar transplantation of shCtrl and shS6K1 2^nd^ tNES in NSG pups at 3-6 days of age, and tumor growth was monitored by IVIS imaging. The IVIS images showed tumor onset at week 7 after transplantation of 2^nd^ tNES shCtrl cells and the signal gradually increased over time. In contrast, no tumor bioluminescence signal was observed in mice injected with 2^nd^ tNES shS6K1 cells (Fig. 7G) and knock down of S6K1 significantly prolonged survival (Fig. 7H), indicating that S6K1 is critical for MB growth *in vivo* and may represent an attractive therapeutic target.

**Figure 7.**
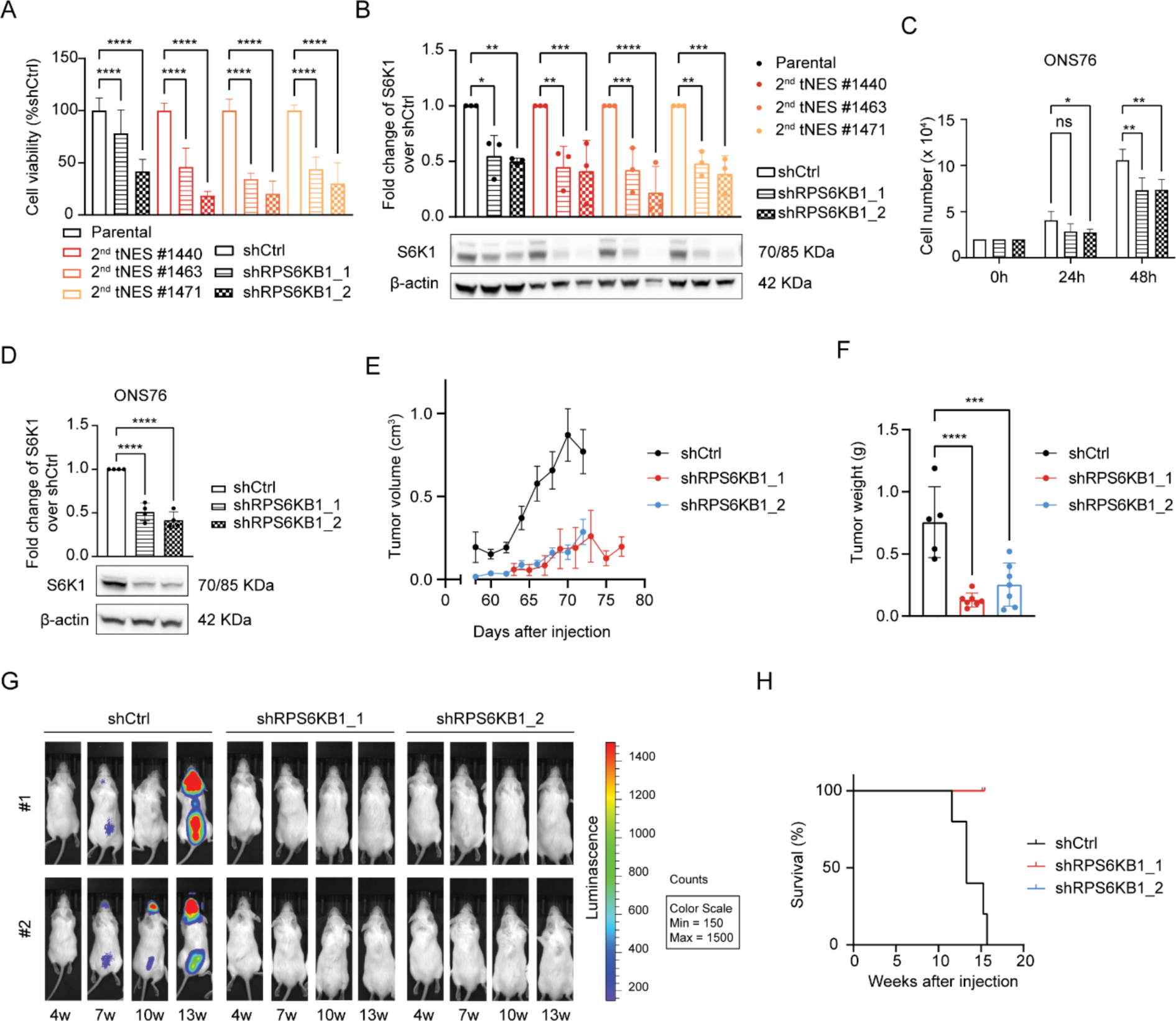
RPS6KB1 knock down decrease tumor cell viability and tumor growth. (A) Cell viability of Parental NES and 2^nd^ tNES (RPS6KB1 WT vs RPS6KB1 KD, n = 3 independent experiments). (B) Western blot and quantification of S6K1 in Parental NES and 2^nd^ tNES (WT vs RPS6KB1 KD, n = 3 independent experiments). (C) Cell proliferation of ONS76 cells (WT vs RPS6KB1 KD, n = 3 independent experiments). (D) Western blot and quantification of S6K1 in ONS76 cells (RPS6KB1 WT vs RPS6KB1 KD, n = 3 independent experiments). (E) Tumor growth of RPS6KB1 WT and RPS6KB1 KD upon subcutaneous injection (n=5 mice in each group). (F) Tumor weight of RPS6KB1 WT and RPS6KB1 KD upon subcutaneous injection at sacrifice (n=5 mice in each group). (G) IVIS imaging of tumor bioluminescence signal reflects tumor growth by 2^nd^ tNES of RPS6KB1 WT and KD. (H) Kaplan-Meier curve reflects survival of mice transplanted with 2^nd^ tNES RPS6KB1 WT and KD (n=5 mice per group).

## Discussion

Although the overall survival of MB patients has increased with combination treatment of surgery, radiation, and chemotherapy, treatment-related side effects and late complications are closely associated with quality of survival. Surgical removal of tumors covering the brainstem or cranial nerves causes posterior fossa syndrome in 25% of patients with MB resection [1]. High doses of irradiation are associated with neurocognitive disability, neuroendocrine dysfunction, growth disturbances, and deformities, which mainly affect younger children. Addition of chemotherapy results in an increase of treatment-related toxicities such as hearing loss, bone marrow aplasia, risk of infection and neurotoxicity [54]. Targeted therapies currently in preclinical and clinical studies are also found to have various adverse effects. SMO inhibitors (Vismodegib and Sonidegib) causes growth plate fusion [21, 55]. Hyperglycemia is commonly seen with PI3K inhibitors [56, 57]. The main adverse effect of CDK inhibitors (Palbociclib, Ribociclib, and Abemaciclib) is leukopenia and neutropenia [58]. Altogether demonstrating the urgent need to develop additional therapies with improved efficacy and minimized side effect for MB treatment. To address these challenges, we used our established human patient-derived NES model to create an assay platform with high-throughput drug screening capacity for parallel drug response assessment in both healthy and malignant cells from the same patient [25]. This enable us to identify compounds and targets that act on selective vulnerabilities for tumor cells, while sparing normal neural stem cells or neurons.

An interesting finding from the screening is that most of the drugs we tested showed similar efficacy against healthy neural stem cells and tumor cells. The reason could be that SHH-MB originates from granule neural precursors (GNPs) which undergo extensive self-renewal during cerebellar development. Many pathways operating during development of normal stem cells are also involved in proliferation of tumor cells [59]. Thus, interfering with such pathways may inhibit the proliferation and survival of both tumor cells and normal neural stem cells (NSCs), and must be considered given that cerebellar development continues after birth into early childhood [60]. Another finding is that inhibition of the MAPK/ERK pathway was more effective in normal NSCs than in tumor cells and it is consistent with previous studies showing that ERK2 inhibition impairs proliferation and self-renewal of NSCs [61, 62].

In our study Brivanib, a selective VEGFR2 and FGFR1 inhibitor that has shown significant anti-tumor activity in clinical trials [63], was found to have a selective anti-proliferative effect on MB tumor cell monolayers, while it unfortunately had no measurable effect on the size of 3D tumor spheroids. The reason may be that Brivanib does not reach the effective dose due to poor spheroid penetration through tight cell-cell adhesion, or that the hypoxic core increase drug resistance mechanisms [64, 65].

In contrast PF4708671, a selective S6K1 inhibitor, showed selective cytotoxicity in 2^nd^ tNES and MB cell lines and effect on the size of 3D tumor spheroids. Moreover, silencing of S6K1 blocked tumor growth *in vivo*. S6K1, a downstream effector of the PI3K/mTOR pathway, is a serine/threonine kinase encoded by the RPS6KB1 gene and has been identified as an important kinase for mitogen-induced phosphorylation of ribosomal S6 protein (S6) to promote global translation and cell growth [66]. S6K1 can be inhibited by targeting either S6K1 or upstream activators e.g. in the PI3K/mTOR pathway. Intriguingly, the PI3K/mTOR pathway inhibitors tested showed equal efficacy on 2^nd^ tNES and Parental NES, while inhibition of S6K1 showed selectivity towards 2^nd^ tNES. The possible mechanism underlying the increased toxicity of S6K1 inhibition may be that activation of PI3K/mTOR pathway in 2^nd^ tNES could sensitize it to S6K1 inhibition. As S6K1 is not the only substrate of the PI3K/mTOR pathway, targeting PI3K or mTORC1 could have a broader effect and reduce the selectivity toward tumor cells. On the other hand, SHH signaling and PI3K/mTOR signaling converge on S6K1 and promote GLI1 oncogenic activity through S6K1-mediated GLI1 phosphorylation [67, 68], suggesting that inhibition of S6K1 blocks tumorigenesis driven by SHH signaling. However, the function of S6K1 in the crosstalk between SHH and PI3K/mTOR pathway in MB needs further investigation.

In the present study, we tested a limited set of biological active molecules, but it will be possible and interesting to expand to significantly more compounds. Also, here we focused specifically on SHH-MB, which represents 30% of MB patients and the effectiveness of the drugs we tested in tNES cells with *PTCH1* mutations might vary in tumors with different genetic changes. Therefore, it will be important to develop and test NES models with various genetic alterations to get a broader understanding of drug sensitivity in MB, stratify patients and develop perhaps precision medicine approaches.

In conclusion, developing safer, more efficient therapies for MB is an urgent need. We show here that patient-derived NES cells can be used for high-throughput screening to identify potential targets for SHH-MB. By targeting specific vulnerabilities, like S6K1, we can hope for improved treatment outcomes and reduced side effects. As we move forward, focusing on refined models and understanding the intricate interplay of signaling pathways will be pivotal for improving MB treatment.

## Data availability

The DiSCoVER method is publicly available as an analysis module in GenePattern (https://www.genepattern.org). Bulk RNA sequencing data of Parental NES and 2^nd^ tNES and microarray expression data of SHH-MB patient and normal cerebellum and upper rhombic lip has been previously published [25, 28] and is available at Gene Expression Omnibus (GSE106718, GSE124814).

## Reference

1. Northcott, P.A., et al., Medulloblastoma. Nat Rev Dis Primers, 2019. 5(1): p. 11.

2. Jessa, S., et al., Stalled developmental programs at the root of pediatric brain tumors. Nat Genet, 2019. 51(12): p. 1702–1713.

3. Louis, D.N., et al., The 2021 WHO Classification of Tumors of the Central Nervous System: a summary. Neuro Oncol, 2021. 23(8): p. 1231–1251.

4. Wechsler-Reya, R.J. and M.P. Scott, Control of neuronal precursor proliferation in the cerebellum by Sonic Hedgehog. Neuron, 1999. 22(1): p. 103–14.

5. Hatten, M.E. and M.F. Roussel, Development and cancer of the cerebellum. Trends Neurosci, 2011. 34(3): p. 134–42.

6. Kool, M., et al., Molecular subgroups of medulloblastoma: an international meta-analysis of transcriptome, genetic aberrations, and clinical data of WNT, SHH, Group 3, and Group 4 medulloblastomas. Acta Neuropathol, 2012. 123(4): p. 473–84.

7. Taylor, M.D., et al., Molecular subgroups of medulloblastoma: the current consensus. Acta Neuropathol, 2012. 123(4): p. 465–72.

8. Ellison, D.W., Childhood medulloblastoma: novel approaches to the classification of a heterogeneous disease. Acta Neuropathol, 2010. 120(3): p. 305–16.

9. Garcia-Lopez, J., et al., Deconstructing Sonic Hedgehog Medulloblastoma: Molecular Subtypes, Drivers, and Beyond. Trends Genet, 2021. 37(3): p. 235–250.

10. Cavalli, F.M.G., et al., Intertumoral Heterogeneity within Medulloblastoma Subgroups. Cancer Cell, 2017. 31(6): p. 737–754.e6.

11. Kool, M., et al., Genome sequencing of SHH medulloblastoma predicts genotype-related response to smoothened inhibition. Cancer Cell, 2014. 25(3): p. 393–405.

12. Schwalbe, E.C., et al., Novel molecular subgroups for clinical classification and outcome prediction in childhood medulloblastoma: a cohort study. Lancet Oncol, 2017. 18(7): p. 958–971.

13. Zhukova, N., et al., Subgroup-specific prognostic implications of TP53 mutation in medulloblastoma. J Clin Oncol, 2013. 31(23): p. 2927–35.

14. Szalontay, L. and Y. Khakoo, Medulloblastoma: an Old Diagnosis with New Promises. Curr Oncol Rep, 2020. 22(9): p. 90.

15. Northcott, P.A., et al., Medulloblastomics: the end of the beginning. Nat Rev Cancer, 2012. 12(12): p. 818–34.

16. Liu, X., et al., Medulloblastoma: Molecular understanding, treatment evolution, and new developments. Pharmacol Ther, 2020. 210: p. 107516.

17. Dlugosz, A., S. Agrawal, and P. Kirkpatrick, Vismodegib. Nat Rev Drug Discov, 2012. 11(6): p. 437–8.

18. Burness, C.B., Sonidegib: First Global Approval. Drugs, 2015. 75(13): p. 1559–66.

19. Rudin, C.M., et al., Treatment of medulloblastoma with hedgehog pathway inhibitor GDC-0449. N Engl J Med, 2009. 361(12): p. 1173–8.

20. Robinson, G.W., et al., Vismodegib Exerts Targeted Efficacy Against Recurrent Sonic Hedgehog-Subgroup Medulloblastoma: Results From Phase II Pediatric Brain Tumor Consortium Studies PBTC-025B and PBTC-032. J Clin Oncol, 2015. 33(24): p. 2646–54.

21. Robinson, G.W., et al., Irreversible growth plate fusions in children with medulloblastoma treated with a targeted hedgehog pathway inhibitor. Oncotarget, 2017. 8(41): p. 69295–69302.

22. Ventarola, D.J. and D.I. Silverstein, Vismodegib-associated hepatotoxicity: a potential side effect detected in postmarketing surveillance. J Am Acad Dermatol, 2014. 71(2): p. 397–8.

23. Mohan, S.V. and A.L. Chang, Management of Cutaneous and Extracutaneous Side Effects of Smoothened Inhibitor Therapy for Advanced Basal Cell Carcinoma. Clin Cancer Res, 2015. 21(12): p. 2677–83.

24. Yauch, R.L., et al., Smoothened mutation confers resistance to a Hedgehog pathway inhibitor in medulloblastoma. Science, 2009. 326(5952): p. 572–4.

25. Susanto, E., et al., Modeling SHH-driven medulloblastoma with patient iPS cell-derived neural stem cells. Proc Natl Acad Sci U S A, 2020. 117(33): p. 20127–20138.

26. Moriconi, C., et al., INSIDIA: A FIJI Macro Delivering High-Throughput and High-Content Spheroid Invasion Analysis. Biotechnol J, 2017. 12(10).

27. Hanaford, A.R., et al., DiSCoVERing Innovative Therapies for Rare Tumors: Combining Genetically Accurate Disease Models with In Silico Analysis to Identify Novel Therapeutic Targets. Clin Cancer Res, 2016. 22(15): p. 3903–14.

28. Weishaupt, H., et al., Batch-normalization of cerebellar and medulloblastoma gene expression datasets utilizing empirically defined negative control genes. Bioinformatics, 2019. 35(18): p. 3357–3364.

29. Badodi, S., et al., Combination of BMI1 and MAPK/ERK inhibitors is effective in medulloblastoma. Neuro Oncol, 2022. 24(8): p. 1273–1285.

30. Zagozewski, J., et al., Combined MEK and JAK/STAT3 pathway inhibition effectively decreases SHH medulloblastoma tumor progression. Commun Biol, 2022. 5(1): p. 697.

31. Baryawno, N., et al., Small-molecule inhibitors of phosphatidylinositol 3-kinase/Akt signaling inhibit Wnt/beta-catenin pathway cross-talk and suppress medulloblastoma growth. Cancer Res, 2010. 70(1): p. 266–76.

32. Yadav, B., et al., Quantitative scoring of differential drug sensitivity for individually optimized anticancer therapies. Sci Rep, 2014. 4: p. 5193.

33. Bai, R.Y., et al., Effective treatment of diverse medulloblastoma models with mebendazole and its impact on tumor angiogenesis. Neuro Oncol, 2015. 17(4): p. 545–54.

34. Slongo, M.L., et al., Functional VEGF and VEGF receptors are expressed in human medulloblastomas. Neuro Oncol, 2007. 9(4): p. 384–92.

35. Santhana Kumar, K., et al., TGF-β Determines the Pro-migratory Potential of bFGF Signaling in Medulloblastoma. Cell Rep, 2018. 23(13): p. 3798–3812.e8.

36. Hartmann, W., et al., Phosphatidylinositol 3’-kinase/AKT signaling is activated in medulloblastoma cell proliferation and is associated with reduced expression of PTEN. Clin Cancer Res, 2006. 12(10): p. 3019–27.

37. Rao, G., et al., Sonic hedgehog and insulin-like growth factor signaling synergize to induce medulloblastoma formation from nestin-expressing neural progenitors in mice. Oncogene, 2004. 23(36): p. 6156–62.

38. Yang, J., et al., Targeting PI3K in cancer: mechanisms and advances in clinical trials. Mol Cancer, 2019. 18(1): p. 26.

39. Ahmad, R., et al., Targeting MUC1-C inhibits the AKT-S6K1-elF4A pathway regulating TIGAR translation in colorectal cancer. Mol Cancer, 2017. 16(1): p. 33.

40. Shen, H., et al., S6K1 blockade overcomes acquired resistance to EGFR-TKIs in non-small cell lung cancer. Oncogene, 2020. 39(49): p. 7181–7195.

41. Theurillat, J.P., et al., URI is an oncogene amplified in ovarian cancer cells and is required for their survival. Cancer Cell, 2011. 19(3): p. 317–32.

42. Berman, A.Y., et al., ERRα regulates the growth of triple-negative breast cancer cells via S6K1-dependent mechanism. Signal Transduct Target Ther, 2017. 2: p. 17035-.

43. Mo, H., et al., S6K1 amplification confers innate resistance to CDK4/6 inhibitors through activating c-Myc pathway in patients with estrogen receptor-positive breast cancer. Mol Cancer, 2022. 21(1): p. 171.

44. Shahsavani, M., et al., An in vitro model of lissencephaly: expanding the role of DCX during neurogenesis. Mol Psychiatry, 2018. 23(7): p. 1674–1684.

45. Kele, M., et al., Generation of human iPS cell line CTL07-II from human fibroblasts, under defined and xeno-free conditions. Stem Cell Res, 2016. 17(3): p. 474–478.

46. Uhlin, E., et al., Derivation of human iPS cell lines from monozygotic twins in defined and xeno free conditions. Stem Cell Res, 2017. 18: p. 22–25.

47. Ianevski, A., A.K. Giri, and T. Aittokallio, SynergyFinder 3.0: an interactive analysis and consensus interpretation of multi-drug synergies across multiple samples. Nucleic Acids Research, 2022. 50(W1): p. W739–W743.

48. Godfrey, D.I., et al., Unconventional T Cell Targets for Cancer Immunotherapy. Immunity, 2018. 48(3): p. 453–473.

49. Roper, S.J., et al., 3D spheroid models of paediatric SHH medulloblastoma mimic tumour biology, drug response and metastatic dissemination. Sci Rep, 2021. 11(1): p. 4259.

50. Nath, S. and G.R. Devi, Three-dimensional culture systems in cancer research: Focus on tumor spheroid model. Pharmacol Ther, 2016. 163: p. 94–108.

51. Richardson, S., et al., Emergence and maintenance of actionable genetic drivers at medulloblastoma relapse. Neuro Oncol, 2022. 24(1): p. 153–165.

52. Čančer, M., et al., Humanized Stem Cell Models of Pediatric Medulloblastoma Reveal an Oct4/mTOR Axis that Promotes Malignancy. Cell Stem Cell, 2019. 25(6): p. 855–870.e11.

53. Pearce, L.R., et al., Characterization of PF-4708671, a novel and highly specific inhibitor of p70 ribosomal S6 kinase (S6K1). Biochem J, 2010. 431(2): p. 245–55.

54. Lafay-Cousin, L. and C. Dufour, High-Dose Chemotherapy in Children with Newly Diagnosed Medulloblastoma. Cancers (Basel), 2022. 14(3).

55. Kimura, H., J.M. Ng, and T. Curran, Transient inhibition of the Hedgehog pathway in young mice causes permanent defects in bone structure. Cancer Cell, 2008. 13(3): p. 249–60.

56. Bendell, J.C., et al., A First-in-Human Phase 1 Study of LY3023414, an Oral PI3K/mTOR Dual Inhibitor, in Patients with Advanced Cancer. Clin Cancer Res, 2018. 24(14): p. 3253–3262.

57. Sweeney, C.J., et al., Phase Ib/II Study of Enzalutamide with Samotolisib (LY3023414) or Placebo in Patients with Metastatic Castration-Resistant Prostate Cancer. Clin Cancer Res, 2022. 28(11): p. 2237–2247.

58. Finn, R.S., A. Aleshin, and D.J. Slamon, Targeting the cyclin-dependent kinases (CDK) 4/6 in estrogen receptor-positive breast cancers. Breast Cancer Res, 2016. 18(1): p. 17.

59. Shackleton, M., Normal stem cells and cancer stem cells: similar and different. Semin Cancer Biol, 2010. 20(2): p. 85–92.

60. Aldinger, K.A., et al., Spatial and cell type transcriptional landscape of human cerebellar development. Nat Neurosci, 2021. 24(8): p. 1163–1175.

61. Imamura, O., et al., Analysis of extracellular signal-regulated kinase 2 function in neural stem/progenitor cells via nervous system-specific gene disruption. Stem Cells, 2008. 26(12): p. 3247–56.

62. Phoenix, T.N. and S. Temple, Spred1, a negative regulator of Ras-MAPK-ERK, is enriched in CNS germinal zones, dampens NSC proliferation, and maintains ventricular zone structure. Genes Dev, 2010. 24(1): p. 45–56.

63. Llovet, J.M. and V. Hernandez-Gea, Hepatocellular carcinoma: reasons for phase III failure and novel perspectives on trial design. Clin Cancer Res, 2014. 20(8): p. 2072–9.

64. Mehta, G., et al., Opportunities and challenges for use of tumor spheroids as models to test drug delivery and efficacy. J Control Release, 2012. 164(2): p. 192–204.

65. Jing, X., et al., Role of hypoxia in cancer therapy by regulating the tumor microenvironment. Mol Cancer, 2019. 18(1): p. 157.

66. Fenton, T.R. and I.T. Gout, Functions and regulation of the 70kDa ribosomal S6 kinases. Int J Biochem Cell Biol, 2011. 43(1): p. 47–59.

67. Wang, Y., et al., The crosstalk of mTOR/S6K1 and Hedgehog pathways. Cancer Cell, 2012. 21(3): p. 374–87.

68. Filbin, M.G., et al., Coordinate activation of Shh and PI3K signaling in PTEN-deficient glioblastoma: new therapeutic opportunities. Nat Med, 2013. 19(11): p. 1518–23.

