## Supplemental Figures and Methods for "High-throughput neural stem cell-based drug screening identifies S6K1 inhibition as a selective vulnerability in SHH-medulloblastoma"

Fig. S1

A

| Percentage of effective compounds predicted in GDSC dataset | | | |
|---|---|---|---|
| Mechanism of action | Selective for secondary tNES (%) | Selective for patient SHH MB (%) | Selective for both (%) |
| Apoptosis | 37.5 | 25.0 | 12.5 |
| Autophagy | 66.7 | 33.3 | 33.3 |
| Cell cycle/DNA damage | 57.1 | 35.7 | 34.3 |
| Cytoskeleton | 28.6 | 28.6 | 28.6 |
| Epigenetics | 50.0 | 33.3 | 33.3 |
| GPCR/G | 0.0 | 0.0 | 0.0 |
| Hormonal | 100.0 | 50.0 | 50.0 |
| JAK/STAT | 33.3 | 0.0 | 0.0 |
| MAPK/ERK | 69.2 | 53.8 | 46.2 |
| Metabolic Enzyme/Protease | 50.0 | 0.0 | 0.0 |
| NF-kB | 25.0 | 25.0 | 0.0 |
| nRTK | 60.0 | 28.0 | 24.0 |
| PI3K/mTOR | 50.0 | 23.3 | 23.3 |
| PKC | 100.0 | 66.7 | 66.7 |
| Protein stability and degradation | 87.5 | 75.0 | 75.0 |
| RARs | 100.0 | 50.0 | 50.0 |
| RTK | 48.4 | 51.6 | 22.6 |
| SHH/WNT | 100.0 | 83.3 | 83.3 |
| Other | 69.2 | 46.2 | 46.2 |

B

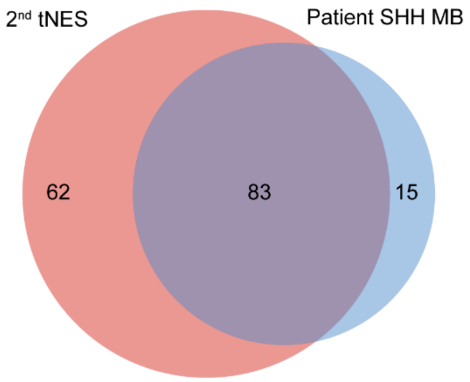


C

| Percentage of effective compounds predicted in CTRPv2 dataset | | | |
|---|---|---|---|
| Mechanism of action | Selective for secondary tNES (%) | Selective for patient SHH MB (%) | Selective for both (%) |
| Apoptosis | 6.9 | 51.7 | 3.4 |
| Autophagy | 33.3 | 55.6 | 22.2 |
| Cell cycle/DNA damage | 18.6 | 66.4 | 15.0 |
| Cytoskeleton | 40.0 | 80.0 | 40.0 |
| Epigenetics | 11.1 | 55.6 | 5.6 |
| GPCR/G | 20.0 | 40.0 | 20.0 |
| Hormonal | 25.0 | 25.0 | 25.0 |
| Immunology/Inflammation | 0.0 | 57.1 | 0.0 |
| JAK/STAT | 30.0 | 100.0 | 30.0 |
| MAPK/ERK | 66.7 | 66.7 | 53.3 |
| Membrane Transporter/Ion Channel | 12.5 | 37.5 | 0.0 |
| Metabolic Enzyme/Protease | 45.0 | 60.0 | 35.0 |
| NF-kB | 28.6 | 57.1 | 28.6 |
| nRTK | 33.3 | 55.6 | 11.1 |
| PI3K/mTOR | 47.8 | 91.3 | 47.8 |
| PKC | 50.0 | 50.0 | 50.0 |
| Protein stability and degradation | 0.0 | 53.3 | 0.0 |
| RARs | 40.0 | 100.0 | 40.0 |
| RTK | 38.6 | 63.6 | 27.3 |
| SHH/WNT | 62.5 | 87.5 | 62.5 |
| Other | 35.8 | 63.3 | 33.3 |

D

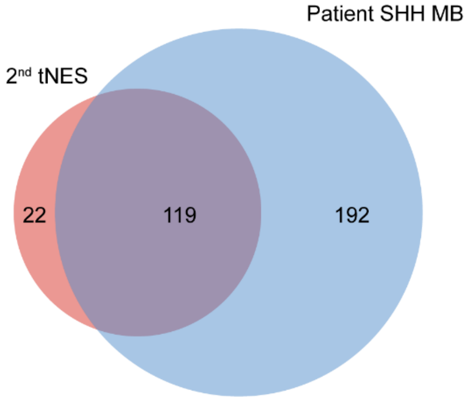

Figure S1. Percentage of compounds predicted efficient against 2$^{nd}$ tNES and patient SHH MB by DiSCoVER. (A) General table of percentage of compounds predicted efficient to 2$^{nd}$ tNES and patient SHH MB from GDSC dataset. (B) Venn diagram showing overlapped effective compounds from GDSC dataset between 2$^{nd}$ tNES and patient SHH MB. (C) General table of percentage of compounds predicted efficient to 2$^{nd}$ tNES and patient SHH MB from CTRPv2 dataset. (B) Venn diagram showing overlapped effective compounds from CTRPv2 dataset between 2$^{nd}$ tNES and patient SHH MB.

Fig. S2

A

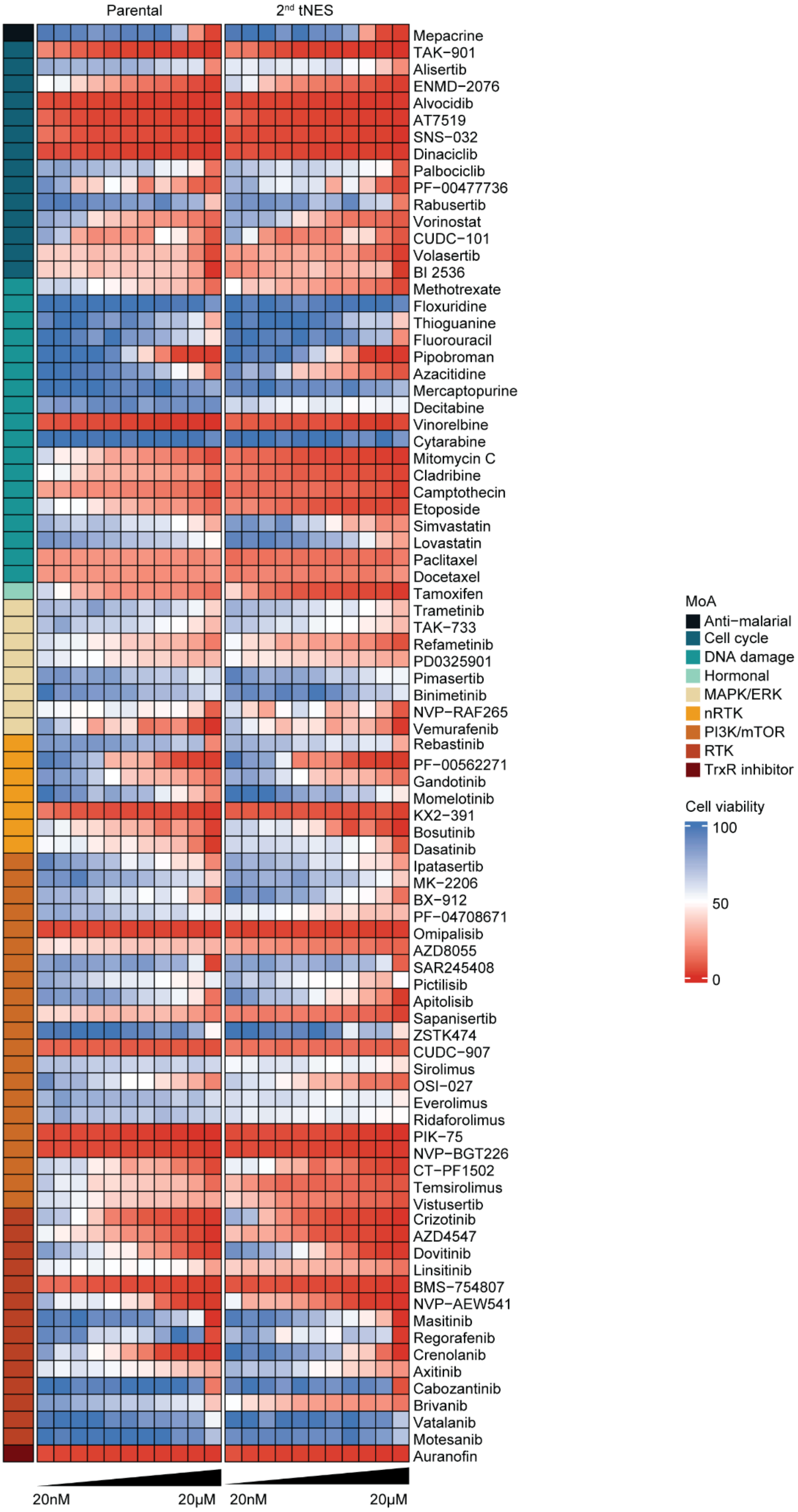
Parental
2nd tNES
Mepacrine
TAK−901
Alisertib
ENMD−2076
Alvocidib
AT7519
SNS−032
Dinaciclib
Palbociclib
PF−00477736
Rabusertib
Vorinostat
CUDC−101
Volasertib
BI 2536
Methotrexate
Floxuridine
Thioguanine
Fluorouracil
Pipobroman
Azacitidine
Mercaptopurine
Decitabine
Vinorelbine
Cytarabine
Mitomycin C
Cladribine
Camptothecin
Etoposide
Simvastatin
Lovastatin
Paclitaxel
Docetaxel
Tamoxifen
Trametinib
TAK−733
Refametinib
PD0325901
Pimasertib
Binimetinib
NVP−RAF265
Vemurafenib
Rebastinib
PF−00562271
Gandotinib
Momelotinib
KX2−391
Bosutinib
Dasatinib
Ipatasertib
MK−2206
BX−912
PF−04708671
Omipalisib
AZD8055
SAR245408
Pictilisib
Apitolisib
Sapanisertib
ZSTK474
CUDC−907
Sirolimus
OSI−027
Everolimus
Ridaforolimus
PIK−75
NVP−BGT226
CT−PF1502
Temsirolimus
Vistusertib
Crizotinib
AZD4547
Dovitinib
Linsitinib
BMS−754807
NVP−AEW541
Masitinib
Regorafenib
Crenolanib
Axitinib
Cabozantinib
Brivanib
Vatalanib
Motesanib
Auranofin
20nM
20μM
20nM
20μM
MoA
Anti−malarial
Cell cycle
DNA damage
Hormonal
MAPK/ERK
nRTK
PI3K/mTOR
RTK
TrxR inhibitor
Cell viability
100
50
0

Figure S2. 85 compounds were tested in 11-point dose-response (20nM-20μM) in singlet to identify compounds showing selectivity towards 2nd tNES. (A) Heatmap showing cell viability of 85 compounds of Parental NES and 2nd tNES.

Fig. S3

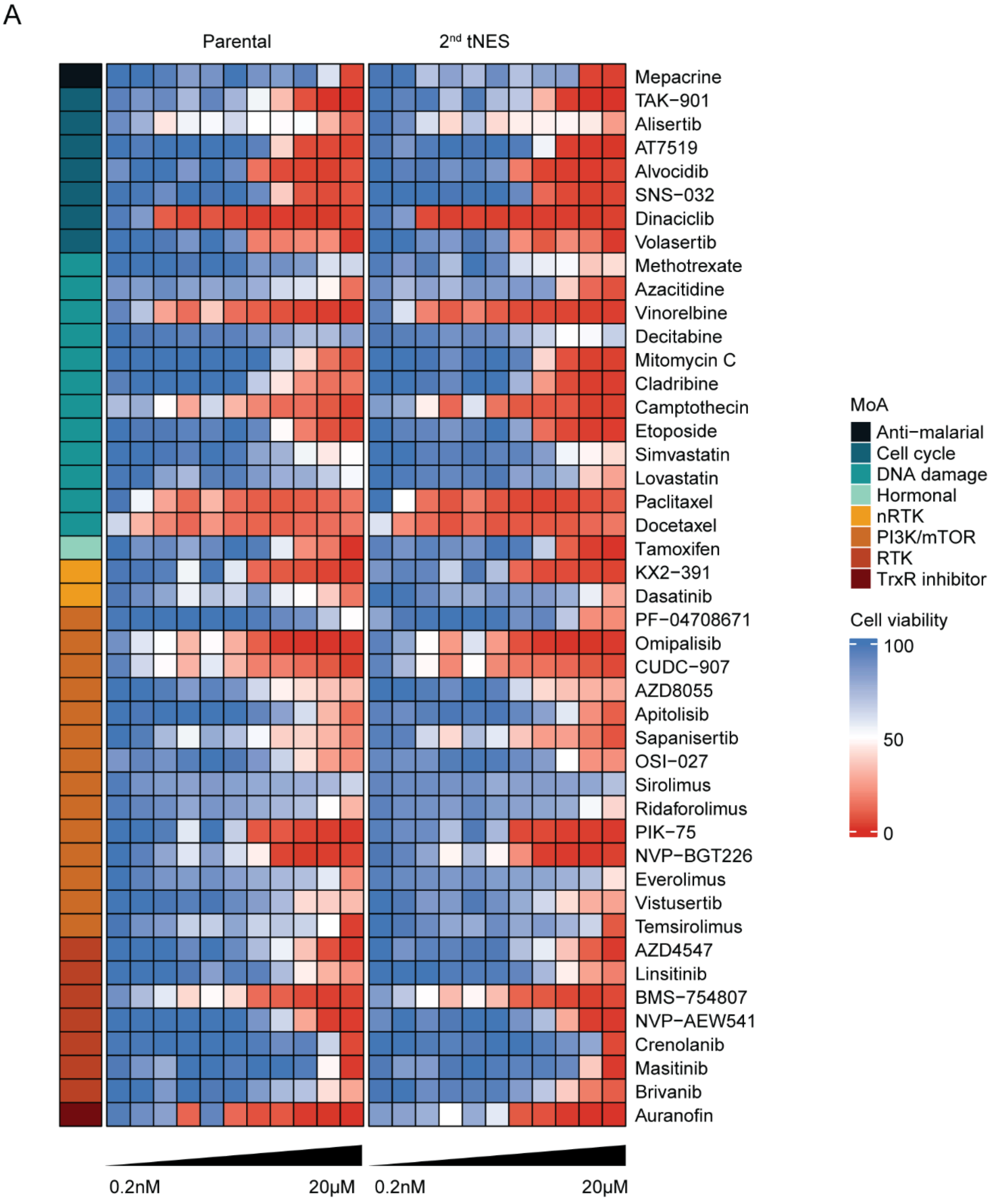
A
Parental
2nd tNES
Mepacrine
TAK−901
Alisertib
AT7519
Alvocidib
SNS−032
Dinaciclib
Volasertib
Methotrexate
Azacitidine
Vinorelbine
Decitabine
Mitomycin C
Cladribine
Camptothecin
Etoposide
Simvastatin
Lovastatin
Paclitaxel
Docetaxel
Tamoxifen
KX2−391
Dasatinib
PF−04708671
Omipalisib
CUDC−907
AZD8055
Apitolisib
Sapanisertib
OSI−027
Sirolimus
Ridaforolimus
PIK−75
NVP−BGT226
Everolimus
Vistusertib
Temsirolimus
AZD4547
Linsitinib
BMS−754807
NVP−AEW541
Crenolanib
Masitinib
Brivanib
Auranofin
0.2nM
20μM
0.2nM
20μM
MoA
Anti−malarial
Cell cycle
DNA damage
Hormonal
nRTK
PI3K/mTOR
RTK
TrxR inhibitor
Cell viability
100
50
0

Figure S3. 46 compounds were tested in 11-point dose-response (0.2nM-20μM) in triplicate to identify compounds showing selectivity towards 2nd tNES. (A) Heatmap showing cell viability of 46 compounds of Parental NES and 2nd tNES.

Fig. S4

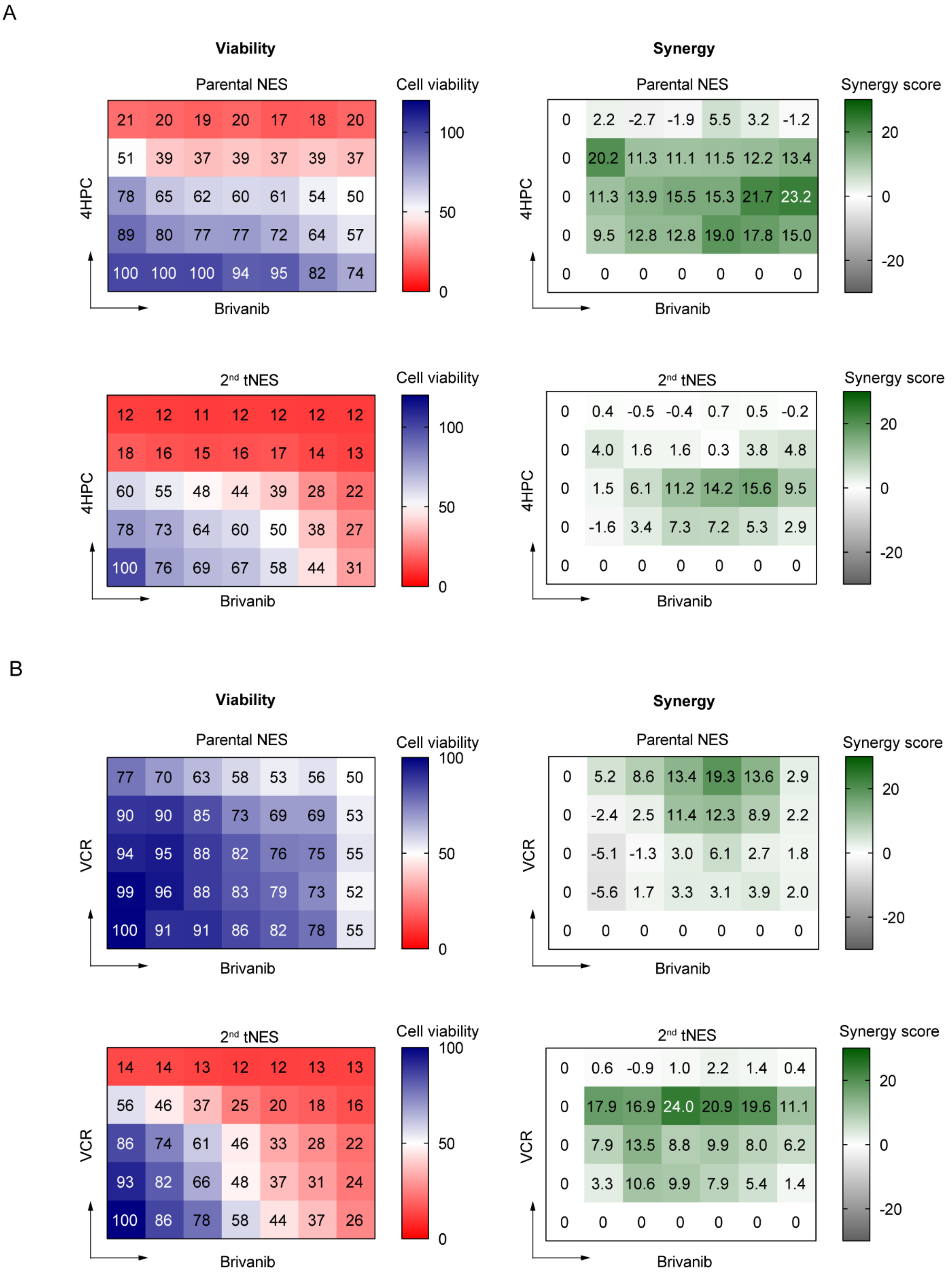
A
Viability
Synergy
Parental NES
Cell viability
100
50
0
21 20 19 20 17 18 20
51 39 37 39 37 39 37
78 65 62 60 61 54 50
89 80 77 77 72 64 57
100 100 100 94 95 82 74
4HPC
Brivanib
Parental NES
Synergy score
20
0
-20
0 2.2 -2.7 -1.9 5.5 3.2 -1.2
0 20.2 11.3 11.1 11.5 12.2 13.4
0 11.3 13.9 15.5 15.3 21.7 23.2
0 9.5 12.8 12.8 19.0 17.8 15.0
0 0 0 0 0 0 0
4HPC
Brivanib
2nd tNES
Cell viability
12 12 11 12 12 12 12
18 16 15 16 17 14 13
60 55 48 44 39 28 22
78 73 64 60 50 38 27
100 76 69 67 58 44 31
4HPC
Brivanib
2nd tNES
Synergy score
0 0.4 -0.5 -0.4 0.7 0.5 -0.2
0 4.0 1.6 1.6 0.3 3.8 4.8
0 1.5 6.1 11.2 14.2 15.6 9.5
0 -1.6 3.4 7.3 7.2 5.3 2.9
0 0 0 0 0 0 0
4HPC
Brivanib
B
Viability
Synergy
Parental NES
Cell viability
77 70 63 58 53 56 50
90 90 85 73 69 69 53
94 95 88 82 76 75 55
99 96 88 83 79 73 52
100 91 91 86 82 78 55
VCR
Brivanib
Parental NES
Synergy score
0 5.2 8.6 13.4 19.3 13.6 2.9
0 -2.4 2.5 11.4 12.3 8.9 2.2
0 -5.1 -1.3 3.0 6.1 2.7 1.8
0 -5.6 1.7 3.3 3.1 3.9 2.0
0 0 0 0 0 0 0
VCR
Brivanib
2nd tNES
Cell viability
14 14 13 12 12 13 13
56 46 37 25 20 18 16
86 74 61 46 33 28 22
93 82 66 48 37 31 24
100 86 78 58 44 37 26
VCR
Brivanib
2nd tNES
Synergy score
0 0.6 -0.9 1.0 2.2 1.4 0.4
0 17.9 16.9 24.0 20.9 19.6 11.1
0 7.9 13.5 8.8 9.9 8.0 6.2
0 3.3 10.6 9.9 7.9 5.4 1.4
0 0 0 0 0 0 0
VCR
Brivanib

Figure S4. Heatmap showing cell viability and compound interactions of different combinations in the Parental NES and $2^{nd}$ tNES. (A) Cell viability of Parental NES and $2^{nd}$ tNES treated with combination of Brivanib and 4HPC (Left), and synergy score of combination of Brivanib and 4HPC (Right). (B) Cell viability of Parental NES and $2^{nd}$ tNES treated with combination of Brivanib and VCR (Left), and synergy score of combination of Brivanib and VCR (Right). All data are from 3 independent experiments.

Fig. S5

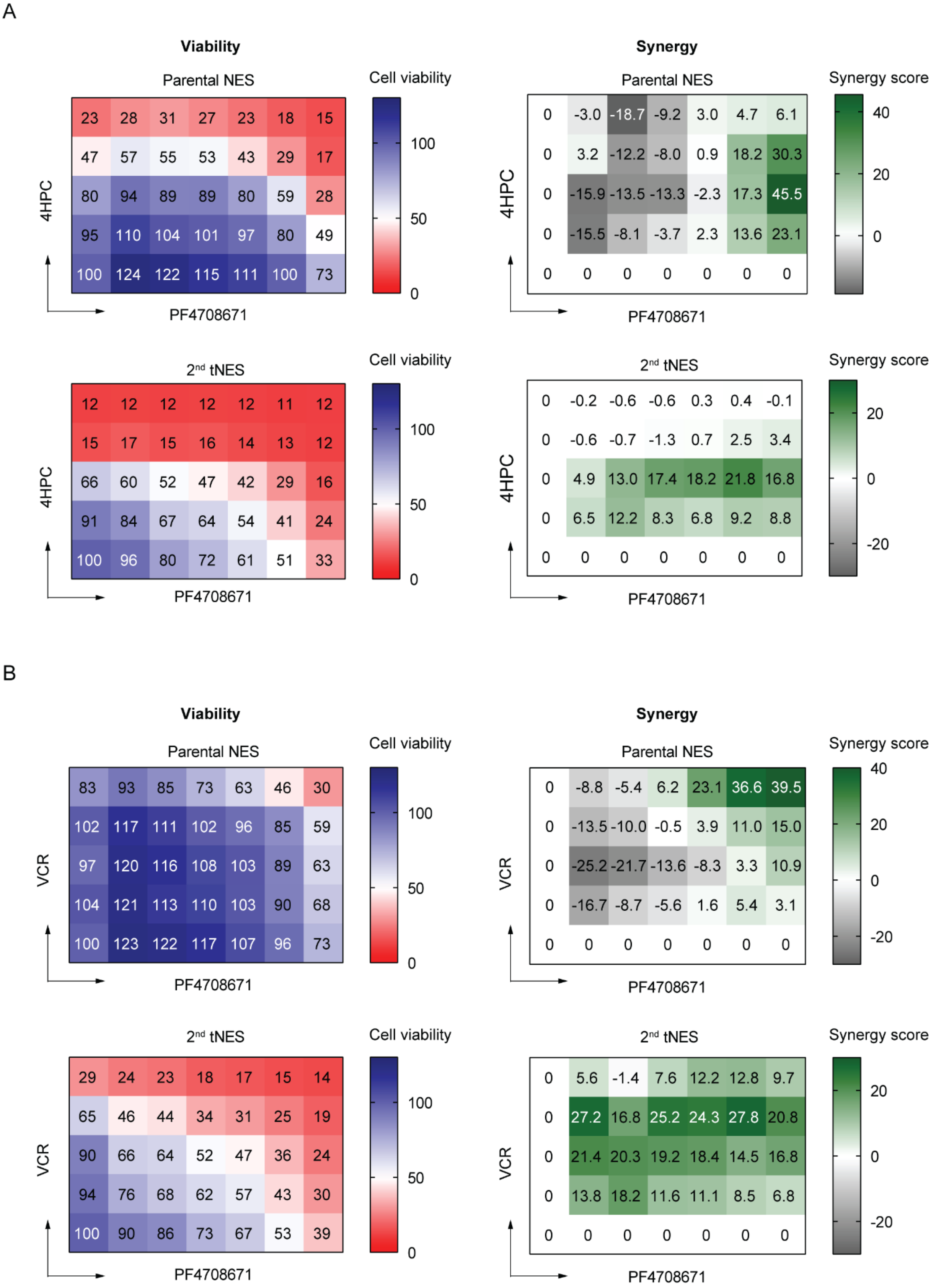
A
Viability
Parental NES
Cell viability
23 28 31 27 23 18 15
47 57 55 53 43 29 17
80 94 89 89 80 59 28
95 110 104 101 97 80 49
100 124 122 115 111 100 73
100
50
0
4HPC
PF4708671
Synergy
Parental NES
Synergy score
0 -3.0 -18.7 -9.2 3.0 4.7 6.1
0 3.2 -12.2 -8.0 0.9 18.2 30.3
0 -15.9 -13.5 -13.3 -2.3 17.3 45.5
0 -15.5 -8.1 -3.7 2.3 13.6 23.1
0 0 0 0 0 0 0
40
20
0
4HPC
PF4708671
2nd tNES
Cell viability
12 12 12 12 12 11 12
15 17 15 16 14 13 12
66 60 52 47 42 29 16
91 84 67 64 54 41 24
100 96 80 72 61 51 33
100
50
0
4HPC
PF4708671
2nd tNES
Synergy score
0 -0.2 -0.6 -0.6 0.3 0.4 -0.1
0 -0.6 -0.7 -1.3 0.7 2.5 3.4
0 4.9 13.0 17.4 18.2 21.8 16.8
0 6.5 12.2 8.3 6.8 9.2 8.8
0 0 0 0 0 0 0
20
0
-20
4HPC
PF4708671
B
Viability
Parental NES
Cell viability
83 93 85 73 63 46 30
102 117 111 102 96 85 59
97 120 116 108 103 89 63
104 121 113 110 103 90 68
100 123 122 117 107 96 73
100
50
0
VCR
PF4708671
Synergy
Parental NES
Synergy score
0 -8.8 -5.4 6.2 23.1 36.6 39.5
0 -13.5 -10.0 -0.5 3.9 11.0 15.0
0 -25.2 -21.7 -13.6 -8.3 3.3 10.9
0 -16.7 -8.7 -5.6 1.6 5.4 3.1
0 0 0 0 0 0 0
40
20
0
-20
VCR
PF4708671
2nd tNES
Cell viability
29 24 23 18 17 15 14
65 46 44 34 31 25 19
90 66 64 52 47 36 24
94 76 68 62 57 43 30
100 90 86 73 67 53 39
100
50
0
VCR
PF4708671
2nd tNES
Synergy score
0 5.6 -1.4 7.6 12.2 12.8 9.7
0 27.2 16.8 25.2 24.3 27.8 20.8
0 21.4 20.3 19.2 18.4 14.5 16.8
0 13.8 18.2 11.6 11.1 8.5 6.8
0 0 0 0 0 0 0
20
0
-20
VCR
PF4708671

Figure S5. Heatmap showing cell viability and compound interactions of different combinations in Parental NES and 2$^{nd}$ tNES. (A) Cell viability of Parental NES and 2$^{nd}$ tNES treated with combination of PF4708671 and 4HPC (Left), and synergy score of combination of PF4708671 and 4HPC (Right). (B) Cell viability of Parental NES and 2$^{nd}$ tNES treated with combination of PF4708671 and VCR (Left), and synergy score of combination of PF4708671 and VCR (Right). All data were from 3 independent experiments.

Fig. S6

A

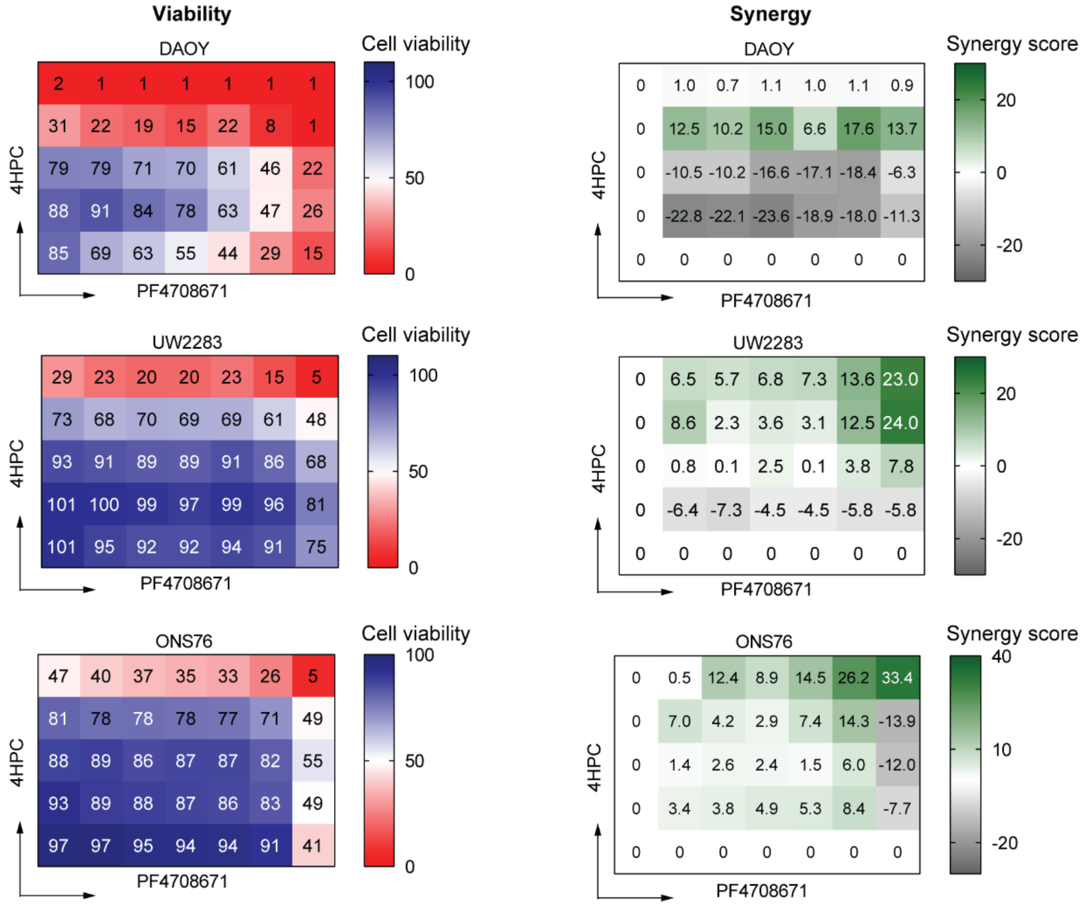

Viability
DAOY
Cell viability
100
50
0
2 1 1 1 1 1 1
31 22 19 15 22 8 1
79 79 71 70 61 46 22
88 91 84 78 63 47 26
85 69 63 55 44 29 15
4HPC
PF4708671
UW2283
Cell viability
100
50
0
29 23 20 20 23 15 5
73 68 70 69 69 61 48
93 91 89 89 91 86 68
101 100 99 97 99 96 81
101 95 92 92 94 91 75
4HPC
PF4708671
ONS76
Cell viability
100
50
0
47 40 37 35 33 26 5
81 78 78 78 77 71 49
88 89 86 87 87 82 55
93 89 88 87 86 83 49
97 97 95 94 94 91 41
4HPC
PF4708671
Synergy
DAOY
Synergy score
20
0
-20
0 1.0 0.7 1.1 1.0 1.1 0.9
0 12.5 10.2 15.0 6.6 17.6 13.7
0 -10.5 -10.2 -16.6 -17.1 -18.4 -6.3
0 -22.8 -22.1 -23.6 -18.9 -18.0 -11.3
0 0 0 0 0 0 0
4HPC
PF4708671
UW2283
Synergy score
20
0
-20
0 6.5 5.7 6.8 7.3 13.6 23.0
0 8.6 2.3 3.6 3.1 12.5 24.0
0 0.8 0.1 2.5 0.1 3.8 7.8
0 -6.4 -7.3 -4.5 -4.5 -5.8 -5.8
0 0 0 0 0 0 0
4HPC
PF4708671
ONS76
Synergy score
40
10
-20
0 0.5 12.4 8.9 14.5 26.2 33.4
0 7.0 4.2 2.9 7.4 14.3 -13.9
0 1.4 2.6 2.4 1.5 6.0 -12.0
0 3.4 3.8 4.9 5.3 8.4 -7.7
0 0 0 0 0 0 0
4HPC
PF4708671


B

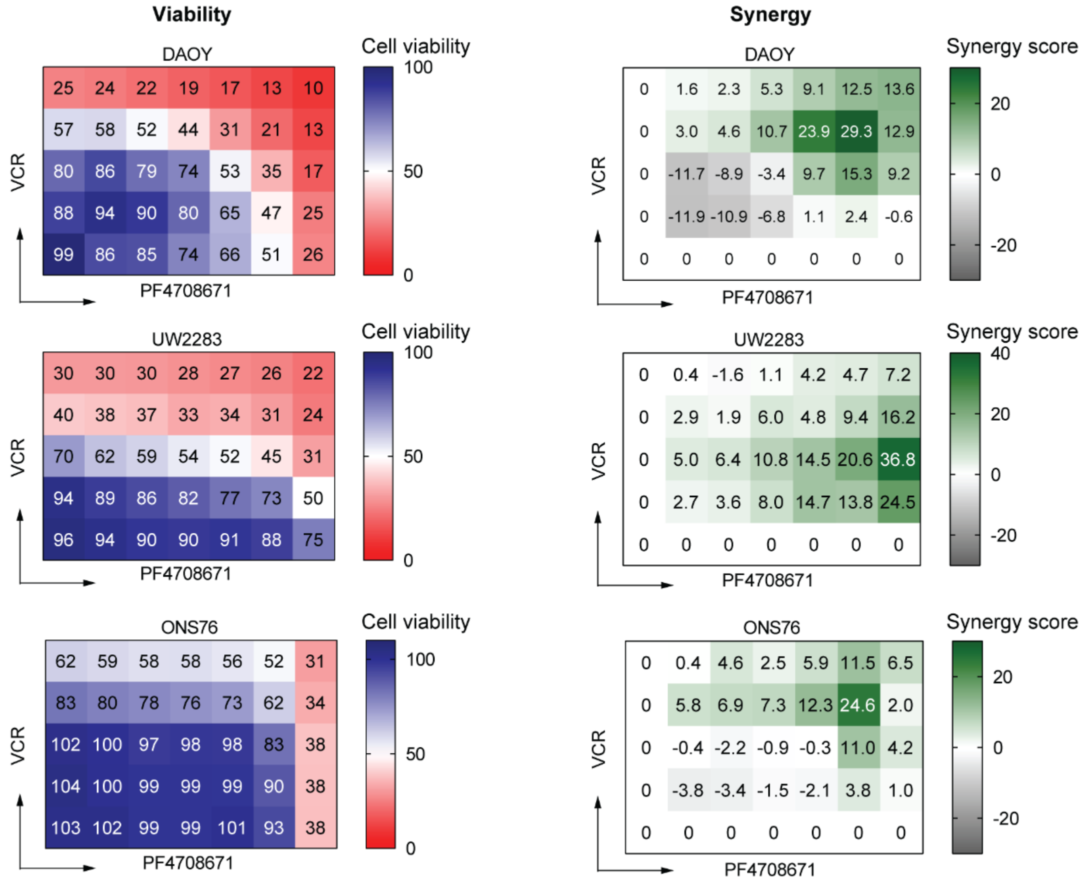

Viability
DAOY
Cell viability
100
50
0
25 24 22 19 17 13 10
57 58 52 44 31 21 13
80 86 79 74 53 35 17
88 94 90 80 65 47 25
99 86 85 74 66 51 26
VCR
PF4708671
UW2283
Cell viability
100
50
0
30 30 30 28 27 26 22
40 38 37 33 34 31 24
70 62 59 54 52 45 31
94 89 86 82 77 73 50
96 94 90 90 91 88 75
VCR
PF4708671
ONS76
Cell viability
100
50
0
62 59 58 58 56 52 31
83 80 78 76 73 62 34
102 100 97 98 98 83 38
104 100 99 99 99 90 38
103 102 99 99 101 93 38
VCR
PF4708671
Synergy
DAOY
Synergy score
20
0
-20
0 1.6 2.3 5.3 9.1 12.5 13.6
0 3.0 4.6 10.7 23.9 29.3 12.9
0 -11.7 -8.9 -3.4 9.7 15.3 9.2
0 -11.9 -10.9 -6.8 1.1 2.4 -0.6
0 0 0 0 0 0 0
VCR
PF4708671
UW2283
Synergy score
40
20
0
-20
0 0.4 -1.6 1.1 4.2 4.7 7.2
0 2.9 1.9 6.0 4.8 9.4 16.2
0 5.0 6.4 10.8 14.5 20.6 36.8
0 2.7 3.6 8.0 14.7 13.8 24.5
0 0 0 0 0 0 0
VCR
PF4708671
ONS76
Synergy score
20
0
-20
0 0.4 4.6 2.5 5.9 11.5 6.5
0 5.8 6.9 7.3 12.3 24.6 2.0
0 -0.4 -2.2 -0.9 -0.3 11.0 4.2
0 -3.8 -3.4 -1.5 -2.1 3.8 1.0
0 0 0 0 0 0 0
VCR
PF4708671

Figure S6. Heatmap showing cell viability and compound interactions of different combinations in DAOY, UW2283 and ONS76 cells. (A) Cell viability of DAOY, UW2283 and ONS76 cells treated with combination of PF4708671 and 4HPC (Left), and synergy score of combination of PF4708671 and 4HPC (Right). (B) Cell viability of DAOY, UW2283 and ONS76 cells treated with combination of PF4708671 and VCR (Left), and synergy score of combination of PF4708671 and VCR (Right). All data were from 3 independent experiments.

Fig. S7

A

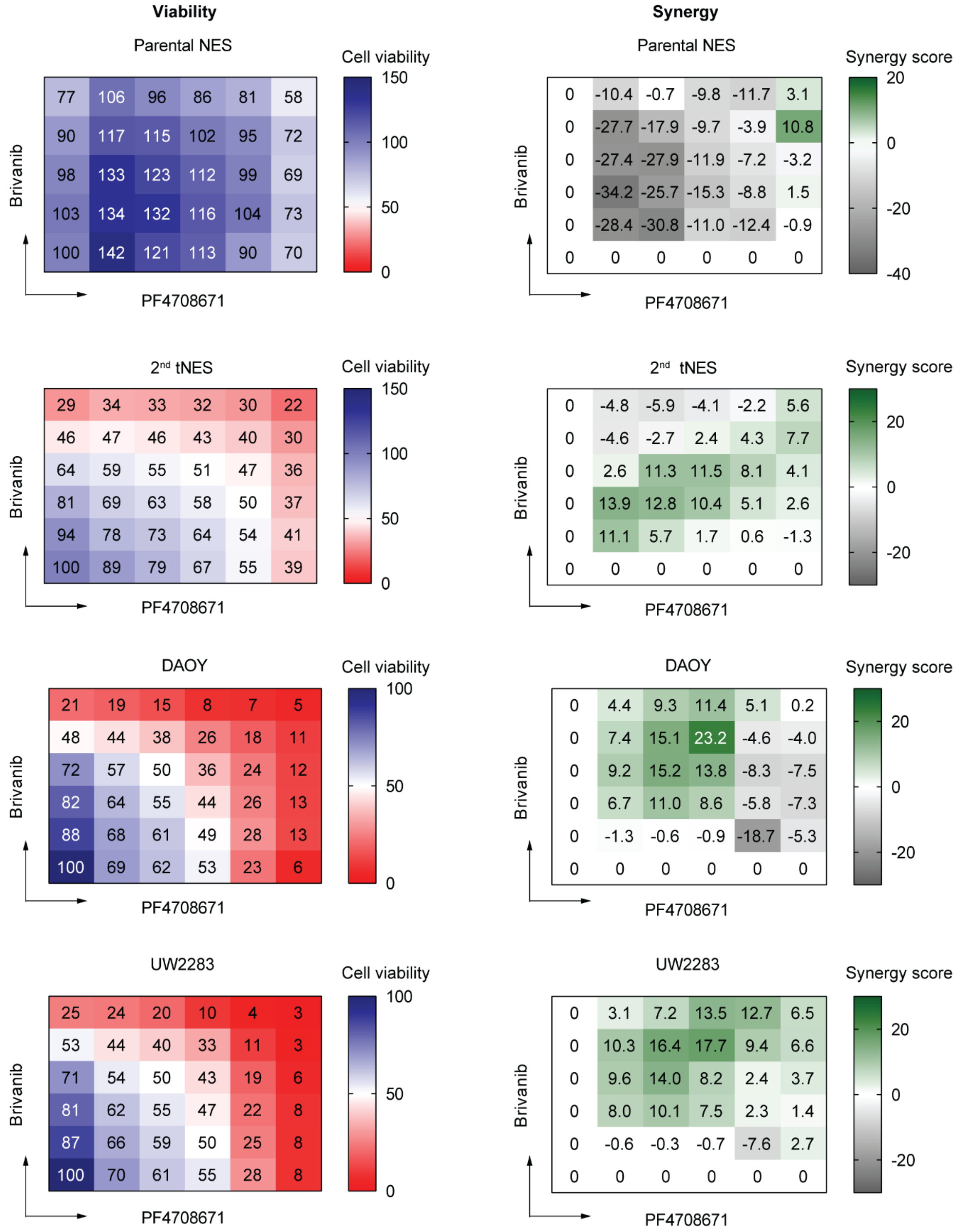
Viability
Parental NES
Cell viability
150
100
50
0
77 106 96 86 81 58
90 117 115 102 95 72
98 133 123 112 99 69
103 134 132 116 104 73
100 142 121 113 90 70
Brivanib
PF4708671
Synergy
Parental NES
Synergy score
20
0
-20
-40
0 -10.4 -0.7 -9.8 -11.7 3.1
0 -27.7 -17.9 -9.7 -3.9 10.8
0 -27.4 -27.9 -11.9 -7.2 -3.2
0 -34.2 -25.7 -15.3 -8.8 1.5
0 -28.4 -30.8 -11.0 -12.4 -0.9
0 0 0 0 0 0
Brivanib
PF4708671
2nd tNES
Cell viability
150
100
50
0
29 34 33 32 30 22
46 47 46 43 40 30
64 59 55 51 47 36
81 69 63 58 50 37
94 78 73 64 54 41
100 89 79 67 55 39
Brivanib
PF4708671
2nd tNES
Synergy score
20
0
-20
0 -4.8 -5.9 -4.1 -2.2 5.6
0 -4.6 -2.7 2.4 4.3 7.7
0 2.6 11.3 11.5 8.1 4.1
0 13.9 12.8 10.4 5.1 2.6
0 11.1 5.7 1.7 0.6 -1.3
0 0 0 0 0 0
Brivanib
PF4708671
DAOY
Cell viability
100
50
0
21 19 15 8 7 5
48 44 38 26 18 11
72 57 50 36 24 12
82 64 55 44 26 13
88 68 61 49 28 13
100 69 62 53 23 6
Brivanib
PF4708671
DAOY
Synergy score
20
0
-20
0 4.4 9.3 11.4 5.1 0.2
0 7.4 15.1 23.2 -4.6 -4.0
0 9.2 15.2 13.8 -8.3 -7.5
0 6.7 11.0 8.6 -5.8 -7.3
0 -1.3 -0.6 -0.9 -18.7 -5.3
0 0 0 0 0 0
Brivanib
PF4708671
UW2283
Cell viability
100
50
0
25 24 20 10 4 3
53 44 40 33 11 3
71 54 50 43 19 6
81 62 55 47 22 8
87 66 59 50 25 8
100 70 61 55 28 8
Brivanib
PF4708671
UW2283
Synergy score
20
0
-20
0 3.1 7.2 13.5 12.7 6.5
0 10.3 16.4 17.7 9.4 6.6
0 9.6 14.0 8.2 2.4 3.7
0 8.0 10.1 7.5 2.3 1.4
0 -0.6 -0.3 -0.7 -7.6 2.7
0 0 0 0 0 0
Brivanib
PF4708671

Figure S7. Heatmap showing cell viability and compound interactions of different combinations in Parental NES, $2^{nd}$ tNES, DAOY and UW2283 cells. (A) Cell viability of Parental NES, $2^{nd}$ tNES, DAOY and UW2283 cells treated with combination of Brivanib and PF4708671 (Left), and synergy score of combination of Brivanib and PF4708671 (Right). All data were from 3 independent experiments.

Fig. S8

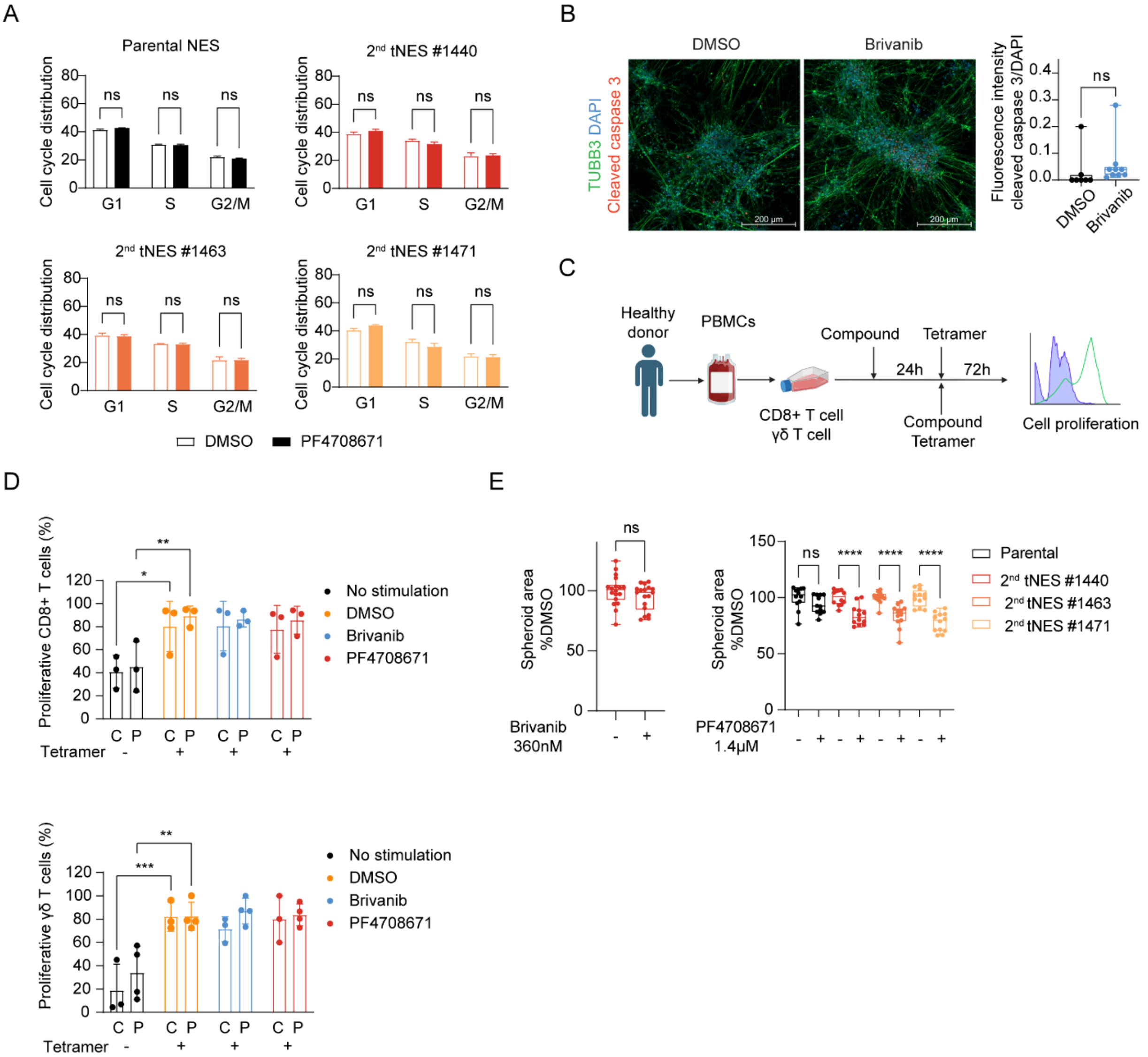
A
Parental NES
2nd tNES #1440
2nd tNES #1463
2nd tNES #1471
Cell cycle distribution
ns
G1
S
G2/M
DMSO
PF4708671
B
DMSO
Brivanib
TUBB3 DAPI
Cleaved caspase 3
200 µm
Fluorescence intensity cleaved caspase 3/DAPI
C
Healthy donor
PBMCs
Compound
Tetramer
24h
72h
CD8+ T cell
γδ T cell
Compound
Tetramer
Cell proliferation
D
Proliferative CD8+ T cells (%)
Proliferative γδ T cells (%)
No stimulation
DMSO
Brivanib
PF4708671
Tetramer
C P
E
Spheroid area %DMSO
Brivanib 360nM
PF4708671 1.4µM
Parental
2nd tNES #1440
2nd tNES #1463
2nd tNES #1471
ns
****

Figure. S8. *In vitro* and *in vivo* effect of Brivanib and PF4708671. (A) Cell cycle analysis of Parental NES and 2$^{nd}$ tNES treated with PF4708671 (n=3 independent experiments). (B) Immunofluorescent staining of beta tubulin III and cleaved caspase 3 in Parental NES differentiated neurons treated with DMSO and Brivanib (360nM) and quantification of cleaved caspase 3. Each dot represents one image. (C) Schematic overview of immune cell proliferation upon compound treatment. (D) Histogram of cell proliferation of CD8+ T and γδ T cell upon Brivanib (360nM) and PF4708671 (750nM) treatment. C indicates combination treatment of compound and tetramer. P indicates prophylactic treatment of compound before addition of tetramer (n=3 independent experiments). (E) Boxplot of size of spheroid derived from 2$^{nd}$ tNES #1440 upon Brivanib (360nM) treatment (n=3 independent experiments) and Parental and 2$^{nd}$ tNES upon PF4708671 (1.4μM) treatment (n=3 independent experiments).

Fig. S9

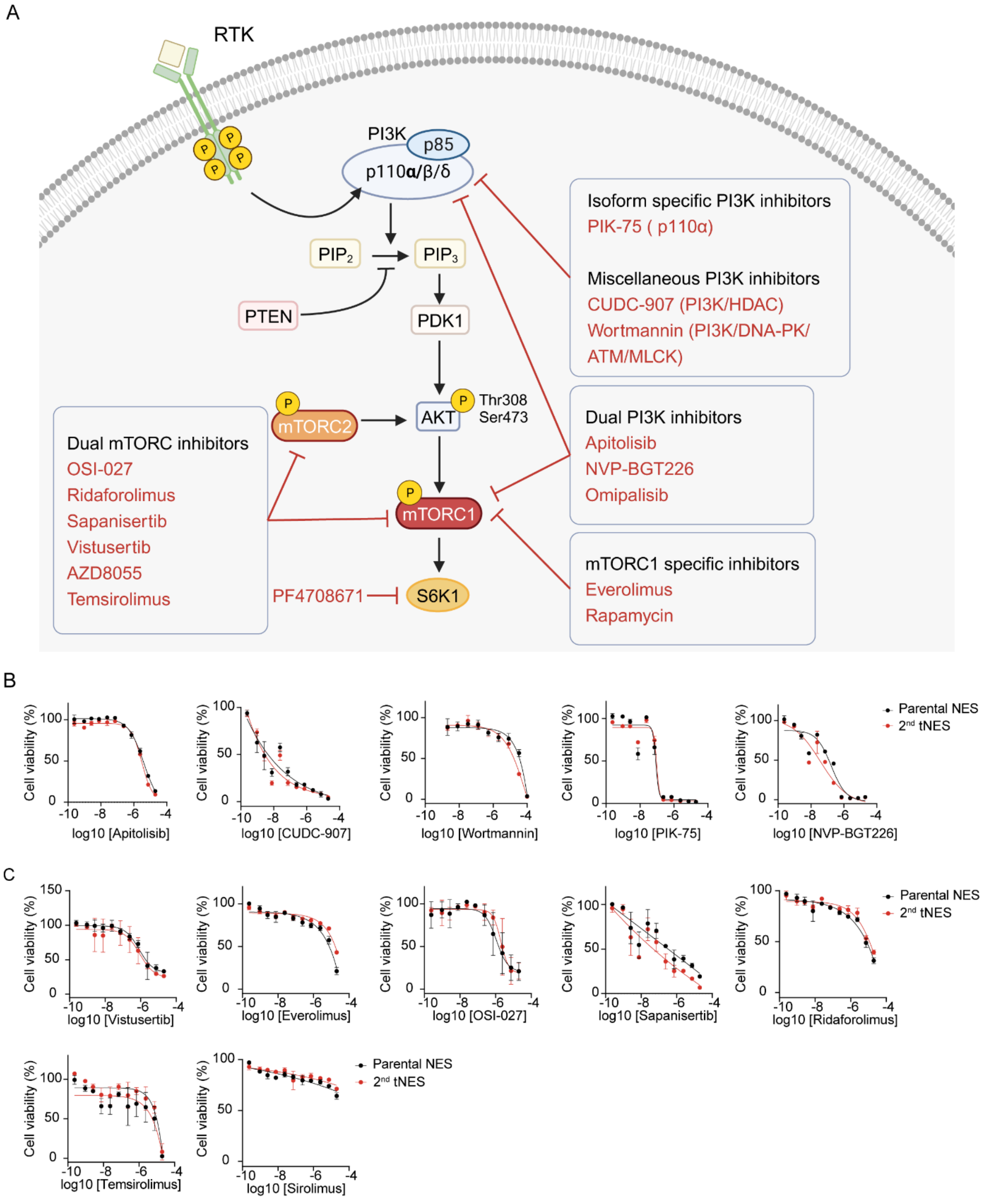
A
RTK
PI3K
p85
p110α/β/δ
PIP2
PIP3
PTEN
PDK1
AKT
Thr308
Ser473
mTORC2
mTORC1
S6K1
PF4708671
Isoform specific PI3K inhibitors
PIK-75 ( p110α)
Miscellaneous PI3K inhibitors
CUDC-907 (PI3K/HDAC)
Wortmannin (PI3K/DNA-PK/ ATM/MLCK)
Dual PI3K inhibitors
Apitolisib
NVP-BGT226
Omipalisib
mTORC1 specific inhibitors
Everolimus
Rapamycin
Dual mTORC inhibitors
OSI-027
Ridaforolimus
Sapanisertib
Vistusertib
AZD8055
Temsirolimus
B
Cell viability (%)
log10 [Apitolisib]
log10 [CUDC-907]
log10 [Wortmannin]
log10 [PIK-75]
log10 [NVP-BGT226]
Parental NES
2nd tNES
C
log10 [Vistusertib]
log10 [Everolimus]
log10 [OSI-027]
log10 [Sapanisertib]
log10 [Ridaforolimus]
log10 [Temsirolimus]
log10 [Sirolimus]

Figure S9. PI3K or mTOR inhibition doesn't result in selectivity towards 2nd tNES. (A) Illustration of PI3K/mTOR pathway showing where the PI3K/mTOR inhibitors target. (B) Dose-response curves of PI3K inhibitors in Parental NES and 2nd tNES. (C) Dose-response curves of mTOR inhibitors in Parental NES and 2nd tNES.

### Materials and Methods

#### Chemicals

Brivanib and PF4708671 were purchased from Selleckchem (Cat#S1084, Cat#S2163). 4-Hydroperoxy Cyclophosphamide was purchased from Santa Cruz Biotechnology (Cat#sc-206885A). Vincristine was a gift from Malin Wickström Lab at Karolinska Institutet. All compound stocks were stored at -80 and aliquots were stored at -20.

#### NES cell lines and culture condition

Generation of Parental NES and 2nd tNES cells was previously described in [1], all NES and tNES cells were grown as monolayers on 0,1 mg Poly-L-ornithine/1µg/ml laminin (Sigma, L2020) coated plates in NES medium containing DMEM/F12 (ThermoFisher Scientific, 31331-028) supplemented with 10% N2 (ThermoFisher, 17504-044), 0.1% B27 (ThermoFisher, 17502-048), 10ng/mL FGF2 (LifeTechnologies, PHG0023) and 10ng/mL EGF (PeproTech, AD100-15). DAOY, UW2283 and HEK293FT cell lines were maintained in DMEM medium containing 1% of Penicillin-Streptomycin (Pen/Strep) (Gibco™, 15070063) and 10% fetal bovine serum (FBS) (Fisher scientific, SV30160.03HI). ONS76 cell line was maintained in RPMI medium (Gibco™, 11875093) containing 1% of Pen/Strep and 10% FBS. Mycoplasma testing was conducted routinely to ensure no contamination. All cells were maintained in a humidified incubator at 37 °C and 5% $CO_2$.

#### DiSCoVER analysis method

The DiSCoVER method is publicly available as an analysis module in GenePattern. The main analysis components of DiSCoVER are shown in Fig 1B. Parental NES and 2nd tNES were profiled using bulk RNA sequencing (GSE106718) and differential gene expression analysis was performed to generate oncogenic signature for 2nd tNES. The enrichment profiles of oncogenic signature, across the cancer cell lines from two mRNA expression dataset, The Cancer Cell Line Encyclopedia (CCLE) and the COSMIC/Sanger Dataset, were computed using single-sample Gene Set Enrichment Analysis (ssGSEA). The enrichment profiles were then matched, using the Information Coefficient (IC) and independently in each dataset, against the sensitivity profiles of The Cancer Therapeutics Response Portal (CTRP v2) and The Genomics of Drug Sensitivity in Cancer (GDSC) over the cell lines where the expression and sensitivity datasets overlap. A matching score of each drug was computed using Information Coefficient (IC) and predicted sensitivity of drugs to 2$^{nd}$ tNES were reflected by the matching scores: a high score indicates likely sensitivity and a low or negative score indicates that the predicted drug was likely to be ineffective [2].

**Cell viability dose-response assay**

Parental NES and 2$^{nd}$ tNES were seeded 15,000 cells/well, Daoy and UW2283 were seeded 5,000 cells/well and ONS76 were seeded 10,000 cells/well in triplicate in 96-well plates the day before start of 48h drug treatment. Cell viability was assessed with Resazurin sodium salt (62.5uM) (Sigma, R7017), which converts to red fluorescent resorufin after incubation with cells for 1-4h and can be measured by a plate reader at excitation/emission 575/585 nm wavelength range. The concentrations of different compounds in different cells were listed in Supplementary Table S4.

**Cytotoxicity assay**

Parental NES and 2$^{nd}$ tNES were seeded at same density as in viability assay in triplicate in the 96-well plate and after 24h they were treated with Brivanib and PF4708671 at EC50 value in 2$^{nd}$ tNES #1440 which is 360nM and 1.4uM, respectively for 48 hours. Cytotoxicity was assessed by CytoTox 96® Non-Radioactive Cytotoxicity Assay kit (Promega, G1780) according to manufacturer's instructions.

**Combination treatment**

Cells were seeded at same density as in single agent viability assay in the 96-well plate for 24h and then treated with combination of Brivanib or PF4708671 and Cyclophosphamide or Vincristine for 48 hours. Cell viability was assessed with Resazurin sodium salt. The expected drug combination responses were calculated based on HSA reference model using SynergyFinder [3]. Deviations between observed and expected responses with positive and negative values denote synergy and antagonism respectively. The concentrations of different compound pairs in different cells are listed in Supplementary Table S5.

**Immune cell proliferation**

Human peripheral blood mononuclear cells (PBMCs) were isolated from blood of healthy donors obtained from Karolinska Hospital (Stockholm, Sweden). PBMCs were collected by Ficoll gradient centrifugation and resuspended in RPMI-1640 medium supplemented with 5% heat-inactivated FBS (Hyclone). T cells were enriched from PBMCs by 30 min-incubation at 37◦C in a horizontal tissue-culture flask (up to 7.106 cells/mL) to promote the adhesion of monocytes, macrophages, and B cells. T cell-enriched supernatants were obtained by centrifugation and labeled with Cell Tracker Yellow (Invitrogen, C34567). DMSO, Brivanib or PF4708671 were either added alone for 24h followed by addition of 25ug/mL tetramer or together with tetramer CD2/CD3/CD28 (Immunocult®, Stemcell) in RPMI-1640 medium supplement with 8% heat-inactivated human serum (Sigma) and 300IU/ml recombinant human IL-2 (StemCell) for 72 hours. Cells were harvested and stained with BV786 conjugated anti-CD3 (BD Biosciences, clone SK7), BV421 conjugated anti-CD8 (BD Bioscience, clone RPA-T8), FITC conjugated anti-pan γδ TCR (Beckman Coulter, clone IMU510) and LIVE/DEAD™ Fixable Aqua (Molecular Probes, L34957) in PBS 0,1%BSA for 30 minutes at 4°C. The proliferation of CD8 positive and γδT cell subsets was measured by flow cytometry using LSRII cytometer (BD Biosicences). Data were analysed using FlowJo software (Treestar).

**Western blot**

For protein detection, cells were washed with PBS and lysed in RIPA buffer (Sigma, R0278) containing 1x Halt™ Protease and Phosphatase Inhibitor Cocktail (ThermoFisher, 78442). Protein concentration of the cell lysates was determined with DC™ Protein Assay Kit II (BioRad, 5000112). The lysates were diluted with 4X Bolt™ LDS Sample Buffer (Invitrogen$^{TM}$, B0008) and boiled for 5min at 95 °C. Equal amounts of total protein extracts were separated by SDS-PAGE (Bolt$^{TM}$ 12% Tris-Bis gel) and electro-transferred onto nitrocellulose membrane with Trans-Blot Turbo Transfer system. The membrane was blocked with 5% nonfat milk or Bovine Serum Albumin in Tris-buffered saline (TBS) containing 0.1% Tween-20 (v/v) for 1 h and incubated with primary antibodies (listed in Supplementary Table S6) at 4°C overnight. The appropriate $2^{nd}$ antibody conjugated to horseradish peroxidase was applied at room temperature for 1 hour. Immunoreactive proteins were developed with ECL SuperSignal West Dura Extended Duration Substrate kit (ThermoFisher, 34076). Blots were quantified by scanning densitometry using area integration.

**Immunofluorescence staining**

Non-guided neuronal differentiation was induced in Parental NES by removal of EGF and FGF2 from NES media, which induced decreased cell proliferation and increased cell differentiation indicated by neurite growth. Media was half-changed every 2 days. After 14 days differentiation, neurons were treated with Brivanib (360nM), PF4708671 (1.4uM) or vincristine (40nM) for 48 hours. After treatment, neurons were fixed in 4% paraformaldehyde for 10 minutes and blocked with 5% goat serum and 0.1% Triton in PBS at room temperature for 1 hour. Neurons were incubated with beta III tubulin and cleaved caspase 3 at 4°C overnight. Neurons were then washed with PBS and incubated with 2$^{nd}$ antibodies conjugated to Alexa fluorophores (1:500) in the dark for 1 hour at room temperature. Nuclear staining was performed using DAPI (1:5000, Sigma, D9542) at room temperature for 5 minutes. Mounting was performed using Fluorescent Mounting medium (Agilent technologies, S302380-2). Primary and 2$^{nd}$ antibodies and dilution are listed in Supplementary Table S6.

**Cell cycle analysis**

Parental NES and 2$^{nd}$ tNES were treated with Brivanib (360nM) and PF4708671 (1.4uM) for 24h. After treatment, floating and adherent cells were collected and washed with PBS and fixed with 70% ethanol at -20°C overnight. Fixed cells were incubated with 100 µg/mL RNase A and stained with 50 µg/mL propidium iodide at 4°C for 30 minutes and then analyzed by Calibur II flow cytometer (BD Biosciences). Cell cycle distribution analysis was performed using FlowJo (FlowJo LLC, Ashland, OR)

**Lentiviral shRNA-mediated knockdown of RPS6KB1**

For preparation of lentiviral particles, pLKO.1-shRPS6KB1 (Sigma, TRCN0000003158, TRCN0000003159) or pLKO.1-shCtrl were transfected with packaging plasmids psPAX2 and pMD2G into HEK293FT cells using lipofectamine 3000 according to manufacturer's instruction (Invitrogen, L3000001). psPAX2 was a gift from Didier Trono (Addgene plasmid # 12260; http://n2t.net/addgene:12260; RRID: Addgene_12260). pMD2.G was a gift from Didier Trono (Addgene plasmid # 12259; http://n2t.net/addgene:12259; RRID: Addgene_12259). After 6 hours of transfection, the cell medium was changed to NES medium or RPMI medium containing 1% Pen/Strep and 10% FBS. Supernatant containing lentiviral particles was harvested at 48 and 72 hours and stored at -80°C. For transduction, NES cells were incubated with lentiviral supernatant (1:5-1:20) for 48 hours. Knockdown efficiency was assessed by western blot of S6K1 protein. Cell viability was assessed with Resazurin sodium salt at 48 hours.

**Zebrafish experiment**

Tg(Fli:EGFP) Zebrafish were mated and embryos were collected and incubated in E3 medium at 28°C until they reach the 1k cell stage which is approximately 4 hours post fertilization. About 100-300 3$^{rd}$ tNES cells stained with Vybrant Dil Cell Label (1:200) was injected into zebrafish embryo using a Sutter P1000 needle puller. After transplantation, embryos were incubated in E3 medium at 33°C. At 24 hours after transplantation, embryos were imaged to sort for ones with intracranial tumors and then treated with PF4708671 (3uM) for 48 hours. Tumor cell viability was assessed by bioluminescent signal.

**Mice experiment**

Human medulloblastoma ONS76 xenograft tumors were established by subcutaneously injection of 2 x $10^6$ cells both RPS6KB1 WT and RPS6KB1 KD suspended in 100uL RPMI medium with 25% Cultrex$^{TM}$ extracellular Matrices into the right flank of NSG mice around 7-8 weeks old. Tumors were measured with calipers every week in the early time and every 2 days when tumor mass observed beneath the skin.

2$^{nd}$ tNES derived tumors were established by orthotopically injection of 50,000 cells both RPS6KB1 WT and RPS6KB1 KD suspended in 1uL NES medium into cerebellum of NSG pups around postnatal day 3-6. Cells were injected using a 30G Hamilton syringe connected to an infusion pump at a rate of 2uL/10 minutes. Needles can penetrate directly into cerebellum hemisphere which is visible at injection. To obtain bioluminescence images, mice were imaged at 10 seconds exposure using IVIS SpectrumCT In Vivo Imaging System (PerkinElmer) within 20 minutes after intraperitoneal injection of D-luciferin (BioThema, BT11-1000K) at dosage of 150mg luciferin/kg body weight.

All mice were fed ad libitum and maintained in environments of 22°C and 12h light and dark cycles. Tumor bearing mice were sacrificed when tumor reached 1.5 $mm^3$ or wound formed on the surface of tumor (s.c) or visible distress. All animal experiments were conducted in accordance with guidelines of Karolinska Institutet and approved by Stockholm's North Ethical Committee of Animal Research (Ethical permits, 6548/18).

1. Susanto, E., et al., *Modeling SHH-driven medulloblastoma with patient iPS cell-derived neural stem cells.* Proc Natl Acad Sci U S A, 2020. **117**(33): p. 20127-20138.
2. Hanaford, A.R., et al., *DiSCoVERing Innovative Therapies for Rare Tumors: Combining Genetically Accurate Disease Models with In Silico Analysis to Identify Novel Therapeutic Targets.* Clin Cancer Res, 2016. **22**(15): p. 3903-14.
3. Ianevski, A., A.K. Giri, and T. Aittokallio, *SynergyFinder 3.0: an interactive analysis and consensus interpretation of multi-drug synergies across multiple samples.* Nucleic Acids Res, 2022. **50**(W1): p. W739-43.
